# Identification of glycosyltransferases that synthesize pectin heteroglycan block polymers and a structural motif required for their activity

**DOI:** 10.64898/2026.10.09.758098

**Authors:** Clifford T. Okoye, Robert A. Amos, Chin Huang, Pradeepa Jayawardhane, Jiri Vlach, Digantkumar Chapla, Jeffrey Ecker, Parastoo Azadi, Kelley W. Moremen, Debra Mohnen

**Affiliations:** Department of Biochemistry and Molecular Biology, University of Georgia, Athens, GA, USA; Complex Carbohydrate Research Center, University of Georgia, Athens, GA, USA

## Abstract

Pectin is a family of plant cell wall complex polysaccharides and proteoglycans comprised of the glycan domains homogalacturonan (HG), rhamnogalacturonan I (RG-I), and the substituted HGs rhamnogalacturonan II (RG-II), xylogalacturonan and apiogalacturonan. The number, type, and arrangement of the pectic domains in different pectic polymers are unknown, as is the identity of the enzymes that link the domains together. Here we show that two Arabidopsis α-1,4-galacturonosyltransferases, GAUT13 and GAUT14, in addition to previously reported HG acceptor-dependent and HG *de novo* synthesis activities, initiate and elongate HG on the non-reducing end of RG-I, identifying enzymes that synthesize pectic RG-I:HG heteroglycans. While other catalytic GAUTs did not initiate HG on RG-I acceptors, they were able to elongate HG once initiated on RG-I by GAUT13 and GAUT14. We show that the three different GAUT14 acceptors (UDP-GalA, HG, and RG-I) compete for the same binding pocket, and amino acids required for GAUT14 catalytic activity were identified using *in vitro* enzyme assays, computational modeling, and site-directed mutagenesis. The transfer of the GAUT14 hypervariable 1 (HV1) motif (RTFASKLQSRS) was sufficient and necessary to transfer RG-I:HG extension activity to GAUT10, which otherwise does not have this activity, representing a novel functional glycosyltransferase domain swapping strategy. The presence of all three catalytic activities in a GAUT13/14 homolog from the non-vascular seedless planst Physcomitrium established the early evolution of GAUT13/14 trifunctionality. The identification of GAUT13 and GAUT14 as RG-I:HG extension catalysts provides targets for probing in planta structure and function of RG-I:HG heteropolymers.

**Teaser:** Two galacturonosyltransferases connect homogalacturonan to rhamnogalacturonan, providing tools to understand and engineer plant cell wall pectic heteroglycan assembly and function.

## INTRODUCTION

Plant cells are surrounded by a dynamic, structurally strong yet malleable extracellular matrix of interacting polymers, including cellulose, hemicellulose, pectin, diverse glycoproteins, and in some cell types, the polyphenolic lignin (*1*). While progress has been made in defining the structures of the major cell wall polysaccharides, less is known about how they are linked together in the wall to form the complex interconnected matrix that imparts cell strength, tissue and organ shape, and enables plant cell growth (*2*). Pectin, a family of galacturonic acid (GalA)-rich polysaccharides, is the most structurally and functionally complex of the cell wall glycans with roles in cell expansion, cell wall structure, cell-to-cell adhesion, pollen tube growth, and plant development (*1, 3, 4*). The most abundant pectin in growing cells is homogalacturonan (HG), a linear glycan of α-1,4-linked D-galacturonic acid (GalA) (*5*). Some portions of HG may be branched with single sugars (xylogalacturonan and apiogalacturonan). Other HG regions are branched with four very complex oligosaccharide side chains and two single sugars yielding the cell wall glycan domain rhamnogalacturonan II (RG-II) (*6*). The other pectic domain, rhamnogalacturonan I (RG-I), is the most abundant pectin in tissues with secondary walls, such as wood, but also constitutes 20-35% of primary wall pectin. RG-I has a repeat backbone of [→4)-α-D-GalA-(1,2)-α-L-Rha-(1→] that is partially branched with neutral sugar side chains including arabinan, galactan, and arabinogalactan (*1, 5, 7, 8*). How the different pectic domains are arranged in the wall is unclear, although it is known that they both covalently and non-covalently associate with each other and with other polymers in the wall (*1, 5*).

The presence of the different pectic domains was identified by their cleavage from the wall using HG hydrolytic enzymes such as endopolygalacturonase (*5*). In this process, information about the structure of unique pectic polymers is lost. The finding that some RG-I and HG in the wall are covalently linked to cell wall arabinogalactan proteins (AGP) (*9, 10*) added a peptide/protein component to pectic structural and functional complexity. More recently, roles for peptides in complex pectin architecture were revealed in the protein-pectin reticulate tripartite tubular lattice network that stabilizes growing pollen tubes, and in the peptide/protein/pectin architecture that compacts HG-containing pectin into periodic circumferential rings essential for root hair expansion (*11, 12*). To understand how HG in such lattices and rings is connected to other pectic polymers and how this influences overall cell wall architecture, it is essential to discern how the different pectic HG and RG-I domains are connected to each other. The goal of this work was to identify enzymes that link the RG-I and HG domains.

Prior chemical analyses of plant cell walls have established that HG exists in the cell wall as a homopolymer (Fig. 1A)(*5*), a heteropolymer with RG-I and HG domains connected via their backbones (Fig. 1B)(*5, 13*), and a heteropolymer with HG and RG-II domains where HG is the backbone of RG-II and extends from both its reducing and non-reducing ends (Fig. 1C)(*14*). HG is also a glycan domain in pectic AGPs, where RG-I is covalently connected to arabinogalactan protein and HG connected to RG-I (Fig. 1D and 1E)(*9, 10*). While the enzymes that synthesize the HG and RG-I backbones have been identified, the enzymes that connect HG to RG-I are not known. HG is synthesized by the GT8 galacturonosyltransferase (GAUT) family (*5, 15*) and the RG-I backbone is synthesized by members of the GT106 RG-I:Rhamnosyltransferase (RRT) family and the GT116 RG-I:Galacturonosyltransferase (RGGAT) family (*16–19*). RRTs and RGGATs work together to synthesize polymeric RG-I by sequential addition of GalA and rhamnose (Rha) on the non-reducing end of growing RG-I oligosaccharides yielding the characteristic RG-I disaccharide repeat backbone (Fig. 1F). Amos et al. (2022) (*19*) have shown that neither RGGATs nor RRTs mediate HG extension of RG-I.

**Fig. 1.**
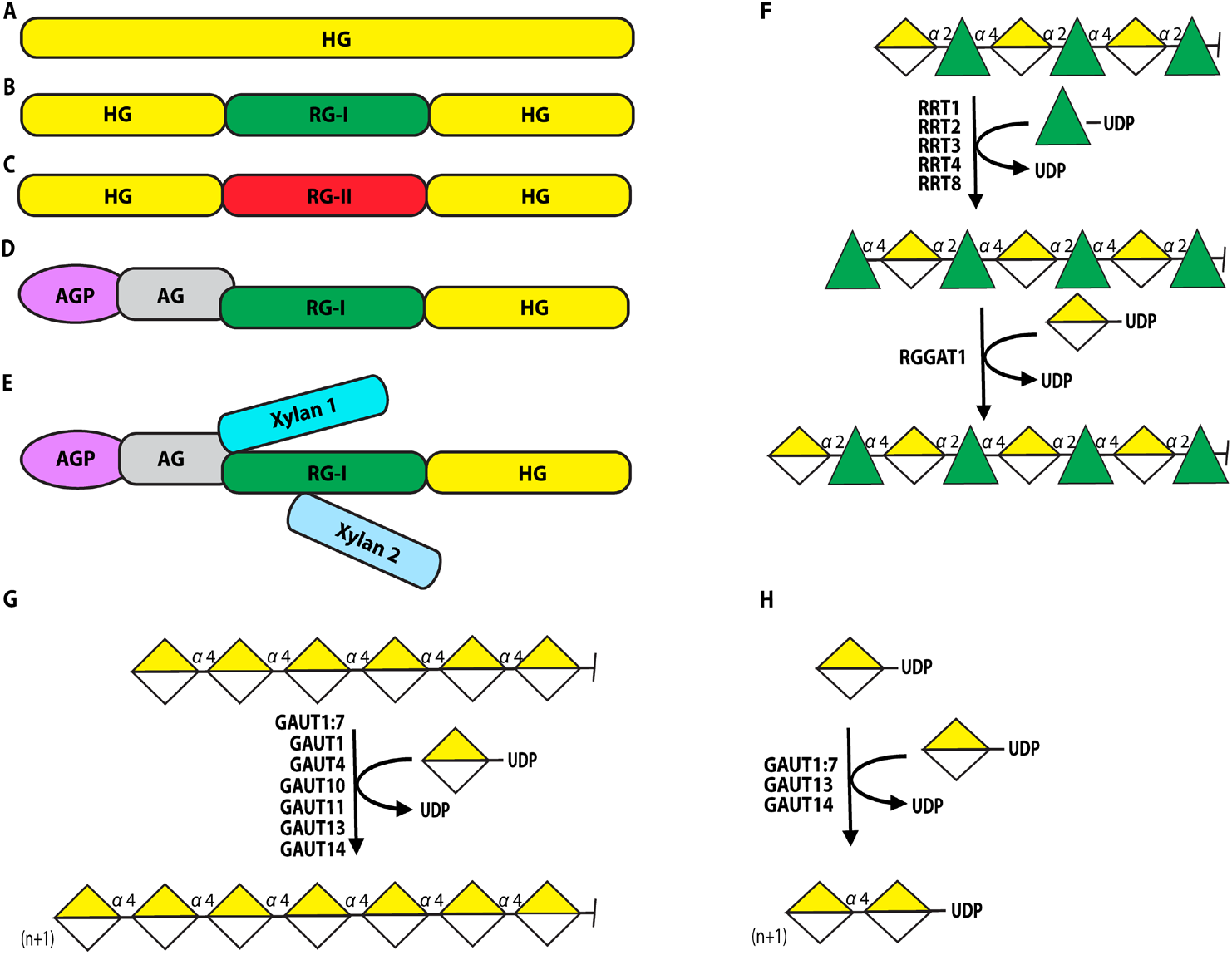
Schematic of documented pectic homoglycans and heteroglycans isolated from the plant cell wall and of enzymes that synthesize the pectic backbones. HG isolated from the plant cell wall exists as either a homopolymer or as HG domains in heteropolymers covalently linked to RG-I or RG-II via the respective backbones. HG domains are present in multiple heteroglycans, including (A) HG homopolymer, (B) RG-I:HG heteropolymer, (C) RG-II:HG heteropolymer, and (D) RG-I:HG domain linked to arabinogalactan proteins (AGP), APAP-1. (E) In Arabidopsis suspension culture medium, the pectic AGP may have attached xylan (*5, 9, 10, 13, 14*). (F) RRTs (RG-I:rhamnosyltransferases) and RGGATs (RG-I:galacturonosyltransferases) catalyze the synthesis of the RG-I backbone by sequential transfer of rhamnose (Rha, green triangle) and galacturonic acid (GalA, yellow and white divided diamond) from their respective nucleotide sugars. (G) All catalytically active GAUTs elongate HG using UDP-GalA as donor substrate and HG oligosaccharides as acceptor substrate (acceptor-dependent activity) (H) Some GAUTs *de novo* synthesize HG using UDP-GalA as both the donor and acceptor substrate (HG *de novo* synthesis activity); (n+1) denotes the addition of one or more GalA residues to the non-reducing end of the HG oligosaccharide.

GAUTs 1, 4, 10, 11, 13, 14, and the GAUT1:GAUT7 complex synthesize α-D-1,4-linked HG through elongation of HG acceptors, an activity called HG acceptor-dependent synthesis (HG:GalAT) (Fig. 1G) (*15, 20–22*). GAUT13, GAUT14, and GAUT1:GAUT7 also have HG *de novo* synthesis activity where they initiate HG synthesis using UDP-GalA as both a donor and an acceptor substrate (Fig. 1H), thereby producing HG product that contains UDP on the reducing end (*15, 21*). No catalytic activity has been demonstrated for GAUTs 5, 6, and 7, however, they have been shown to function as subunits of GAUT1 protein complexes, anchoring catalytic GAUT1 in the Golgi (*20, 23*). All catalytic GAUTs use a retaining glycosyl transfer mechanism to transfer GalA from the donor substrate UDP-α-D-GalA onto the 4-position of HG, generating α-1,4-linked HG. The GalA residue that connects RG-I to HG also has an α-1,4-linkage. Thus, the GAUTs are potential candidate enzymes for initiation of HG synthesis on RG-I acceptors. Although it was previously reported that GAUTs 1, 10, 11, 13, or 14 did not catalyze significant HG synthesis when RG-I was used as an acceptor substrate (*15*), higher-sensitivity enzyme assays and a more complete series of RG-I acceptors are now available to test this more thoroughly.

Here, we show that Arabidopsis GAUT13 and GAUT14 catalyze the addition of GalA from UDP-GalA in an α(1→4)-D-GalA linkage onto the non-reducing end of RG-I backbone acceptors, thereby both initiating and elongating HG on RG-I. We also show that other GAUTs, while not able to initiate HG synthesis on RG-I, are able to elongate the HG once initiated. Using *in vitro* enzyme assays, computational modeling, and site-directed mutagenesis we further identify in GAUT14 amino acids and motifs essential for RG-I:HG extension activity. Finally, we show that RG-I:HG extension activity arose early during plant evolution and is present in the extant non-vascular seedless plant Physcomitrium. The identification of two trifunctional GAUTs with the ability to initiate HG synthesis on RG-I provides targets for the study of pectic heteroglycan function in plants and across plant evolutionary time.

## RESULTS

### GAUT13 and GAUT14 initiate HG synthesis on the non-reducing end of RG-I

The goal of this study was to identify and characterize enzyme(s) that synthesize pectic RG-I:HG heteroglycans. Since multiple members of the GAUT gene family initiate and/or elongate α-D-1,4-linked HG, we hypothesized that one or more GAUTs might catalyze the synthesis of α-D-1,4-linked HG on the non-reducing end of RG-I backbone acceptors. To test this, we expressed and purified all Arabidopsis GAUT proteins previously shown to have HG:GalAT activity when expressed in human embryonic kidney (HEK293F) cells (GAUT1:GAUT7 complex, GAUT10, GAUT11, GAUT13, and GAUT14) (*15, 21*) (Fig. S1A). Transmembrane-domain-truncated GAUT coding regions (GAUT1Δ167:GAUT7Δ43 complex, GAUT10Δ39, GAUT11Δ39, GAUT13Δ61, and GAUT14Δ61) were cloned into pGEn2-DEST (a mammalian expression vector containing a signal peptide sequence, 8×His and Avi tag, and superfolder GFP), heterologously expressed in HEK293F cells, and purified by Ni^2+^-NTA affinity chromatography as previously described (*15, 20, 21, 24*). Recovery of proteins, protein purity, and molecular weight were confirmed by sodium dodecyl sulfate-polyacrylamide gel electrophoresis (SDS-PAGE) under reducing conditions (Fig. S1A). RG-I backbone oligosaccharides with GalA on the non-reducing end (RG-I(G)) were isolated from Arabidopsis seed mucilage for use as acceptor substrates (Fig. S1B-D). These oligosaccharides were derivatized at the reducing end with 2-aminobenzamide (2AB) to yield a fluorescent and UV-absorbing tag, followed by size purification and mass confirmation (see methods, Fig. S1B-D) (*15, 19, 21*).

To test whether any GAUTs could initiate synthesis of HG on the non-reducing end of RG-I acceptors containing a terminal non-reducing GalA residue, 100 µM of purified 2-AB-labeled DP8 RG-I acceptor substrate (RG-I(G)-DP8) was incubated with 1 mM UDP-GalA donor substrate in reactions containing 50 mM HEPES (pH 7.2) and 0.25 mM MnCl_2_ at 30°C, conditions previously shown to be favorable for HG synthesis by the GAUT1:GAUT7 complex (*21*). Enzyme concentrations of 1 µM GAUT1Δ167:GAUT7Δ43 complex, GAUT13Δ61, and GAUT14Δ61, and 10 µM GAUT10Δ39 and GAUT11Δ39 were used since prior reports showed the latter two enzymes to have relatively low HG acceptor-dependent activity (*15*). Products formed after one hour (Fig. S2A) and 12 hours (Fig. 2A) were analyzed by anion-exchange chromatography over an analytical CarboPac^TM^ PA1 column (4 × 250 mm) with fluorescence detection. Peaks eluting later than the RG-I(G)-DP8 acceptor substrate, consistent with the formation of larger-sized charged products, were observed at both 1 hour (Fig. S2A) and 12 hours (Fig. 2A) for GAUT13 and GAUT14. MALDI-TOF MS analysis revealed time-dependent 176 Da increases in the masses of the products, consistent with the stepwise addition of GalA onto the starting RG-I acceptor (Fig. 2B) and (Fig. S2B). The products generated by GAUT13 and GAUT14 in large-scale reactions were further size-purified, and their masses were confirmed by MALDI-TOF MS analysis to correspond to the addition of one, two, three, or four GalA residues to RG-I(G)-DP8, *i.e*., RG-I(G)-DP8 +G, +GG, +GGG, and +GGGG, respectively (Fig. S2C-D). These results demonstrated that GAUT13 and GAUT14 were able to extend the RG-I oligosaccharide acceptor substrates with one to four GalA residues, while no activity was observed for the GAUT1:GAUT7, GAUT10 and GAUT11 enzymes. However, consistent with previous findings by Engel *et al.* (2022)(*15*), these other GAUT enzymes were capable of HG acceptor-dependent activity and HG *de novo* synthesis (Fig. S3A-D)

**Fig. 2.**
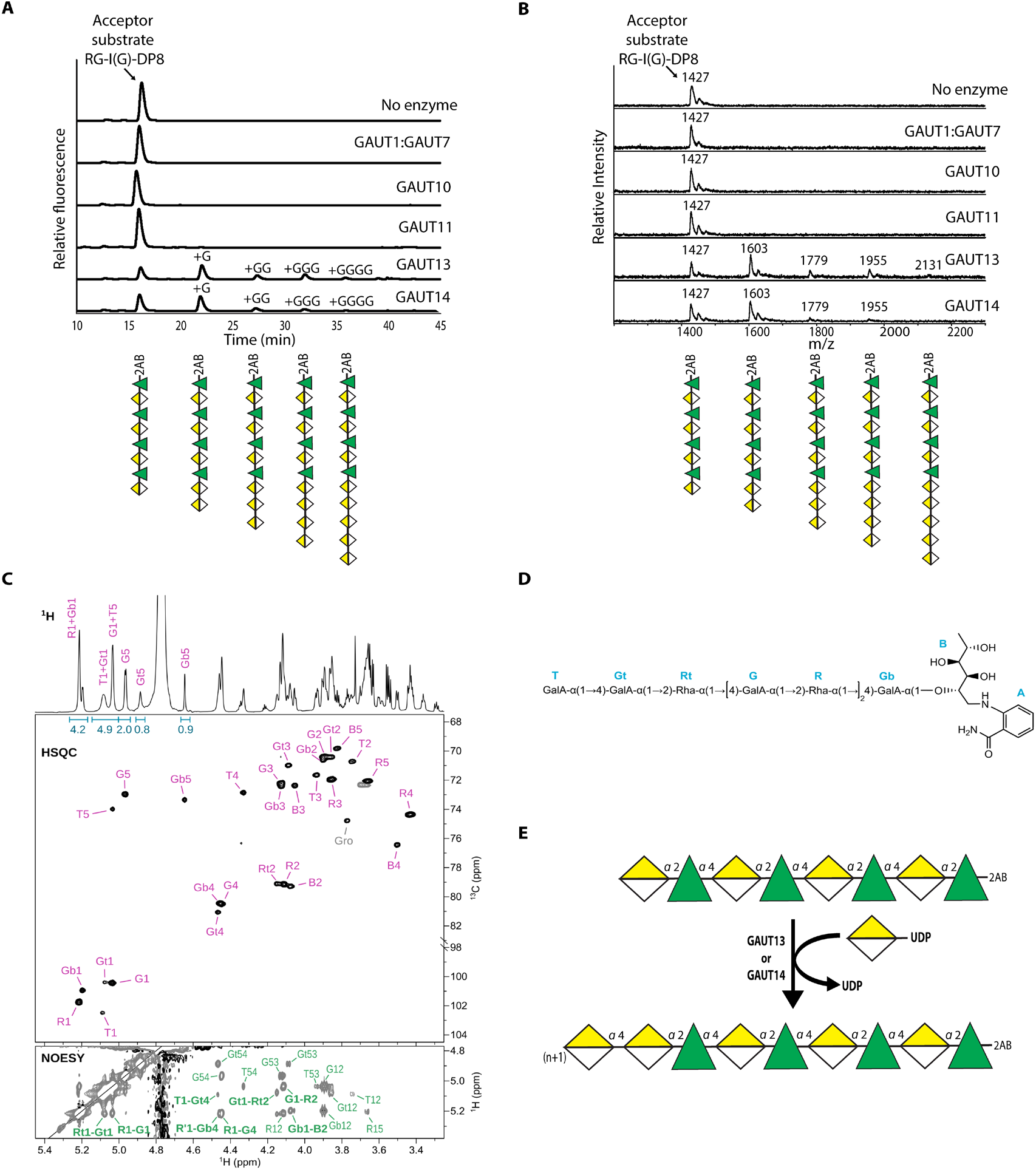
GAUT13 and GAUT14 initiate α(1→4) linked HG on the non-reducing end of RG-I. **(A)** Anion-exchange fluorescence chromatogram of 2AB-labeled reaction products from 12-hour GAUT-catalyzed reactions. Reactions contained 50 mM HEPES (pH 7.2), 0.25 mM MnCl2, 100 µM 2AB-labeled RG-I(G)-DP8 acceptor substrate and 1 mM UDP-GalA donor substrate with the specified enzyme concentration: 1 µM for GAUT1:GAUT7, GAUT13, or GAUT14, or 10 µM for GAUT10 and GAUT11. The difference in enzyme concentration was due to previously reported low HG acceptor-dependent activity for GAUT10 and GAUT11 (*15*). Four 2AB-labeled product peaks are present in chromatograms of GAUT13 and GAUT14 12-hour reactions, eluting between 22 and 37 minutes. The 2AB-labeled RG-I(G)-DP8 acceptor substrate eluted at 16 minutes. Glycan structures of RG-I acceptors and reaction products are depicted according to the Symbol Nomenclature for Glycans (SNFG) (*57*). **(B)** MALDI-TOF mass spectra of reaction products shown in (A), confirming mass of the 2AB-labeled RG-I(G)-DP8 acceptor substrate (mass = 1427 Da) and its elongated products, with mass increases of 176 Da per GalA addition (+G = 1603 Da, +GG = 1779 Da, +GGG = 1955 Da, +GGGG = 2131 Da). The results confirm that only GAUT13 and GAUT14 initiate GalA transfer onto the RG-I(G)-DP8 oligosaccharide acceptor and further elongate that reaction product with additional GalA residues. **(C)** Carbohydrate region of ^1^H (top), ^13^C-HSQC (middle), and NOESY (bottom) spectra of RG-I(G)-DP8+G acquired at pH 2.7. Assignments and integrals for the anomeric H-1 and H-5 signals are indicated in the ^1^H spectrum. Carbohydrate signals are annotated in the HSQC spectrum using the same residue codes as in the chemical shift assignment in Table S1. In the NOESY spectrum, inter-residue signals that identify the glycosidic linkages are shown in boldface. **(D)** Chemical schematic of the 2AB-labeled RG-I(G)-DP8+G oligomer determined from NMR analyses. The terminal and penultimate GalA residues are labeled T and Gt, respectively. Additional residues are annotated as follows: Rt (Rha residue adjacent to the penultimate GalA at the non-reducing end), G and R (internal GalA and Rha residues of the backbone), Gb (GalA adjacent to the reducing-end 1N-rhamnitol), and B (the reducing-end 1N-Rha-ol). The 2-AB fluorophore label at the reducing end is also shown (A). The α(1→4) and α(1→2) glycosidic linkages identified by NOESY and HMBC are indicated. **(E)** Schematic depiction of RG-I:HG extension activity of GAUT13 and GAUT14. (n+1) denotes the addition of one or more GalA residues onto the non-reducing end of the RG-I(G)-DP8 oligosaccharide.

The anomeric configuration and glycosidic linkage of the GalA added onto the non-reducing end of the 2-AB-derivatized RG-I(G) DP8 acceptor were confirmed by NMR spectroscopy of the RG-I(G)-DP8+G extension product. 1D and 2D NMR analysis (Fig. 2C, D, and Table S1) confirmed that a GalA residue was added onto the non-reducing end of the RG-I acceptor and established that the linkage between the terminal and penultimate GalA residues (labeled T and Gt, respectively) was α(1→4) based on NOE correlations and characteristic chemical shifts. These results confirmed that the glycan synthesized on the non-reducing end of the RG-I oligosaccharide acceptor by GAUT13 and GAUT14 had the structure of HG. This activity, illustrated in the reaction scheme shown in Fig. 2E, is named RG-I:HG extension activity.

### The GAUT1:7 Complex, GAUT10, and GAUT11 lack RG-I:HG extension activity but elongate the HG domain once initiated by GAUT13 and GAUT14

The synthesis of biological polymers such as pectin occurs in three stages: initiation, elongation, and termination. Above, we showed that GAUT13 and GAUT14 initiated HG synthesis on the RG-I backbone and that GAUT10, GAUT11, and the GAUT1:GAUT7 complex did not. Once initiated, however, the HG domain must be further elongated to yield the RG-I:HG heteroglycans found in the plant cell wall (*1, 5*). To determine which GAUTs function in the elongation phase of RG-I:HG heteroglycan synthesis, we tested the GAUT1:7 complex, GAUT10, GAUT11, as well as GAUT13 and GAUT14 for the ability to elongate RG-I:HG heteroglycans containing one or more α-1,4-linked GalA residues initiated onto RG-I by GAUT13 and GAUT14 (see Fig. 2A-B).

Size-purified RG-I:HG extension products containing one, two, three, or four α-1,4-linked GalA residues (+G, +GG, +GGG, and +GGGG, respectively) on the non-reducing end of RG-I(G)-DP8 acceptors containing a terminal GalA residue (Fig. S2C-D) were used as acceptors. The 2-AB-labeled reaction products produced by GAUT1:7, GAUT10, GAUT11, GAUT13, and GAUT14 in 1-hour reactions were characterized by anion-exchange chromatography coupled with fluorescence detection. Significantly, while GAUT1:7 did not show detectable GalA transfer to RG-I-DP8 +G acceptor (i.e. RG-I-DP8 with one added α-1,4-linked GalA), GAUT10 and GAUT11 did (Fig. 3A). GAUT1:7, however, was able to transfer GalA to acceptors with two, three or four non-reducing end α-1,4-linked GalA residues (RG-I-DP8 +GG, RG-I-DP8 +GGG, and RG-I-DP8 +GGGG) and more products formed as the acceptor length increased from +GG to +GGG to +GGGG (Fig. 3B-D). GAUT10 and GAUT11, in contrast, elongated all RG-I-DP8:HG acceptors tested, showing preference for RG-I(G)-DP8+GGG (Fig. 3A-D). GAUT13 and GAUT14, in addition to initiating the first α1-4-linked GalA residue onto the RG-I acceptor, also elongated all four RG-I:HG substrates, being most active on RG-I-DP8 +GGGG acceptor (Fig. 3A-D).

**Fig. 3.**
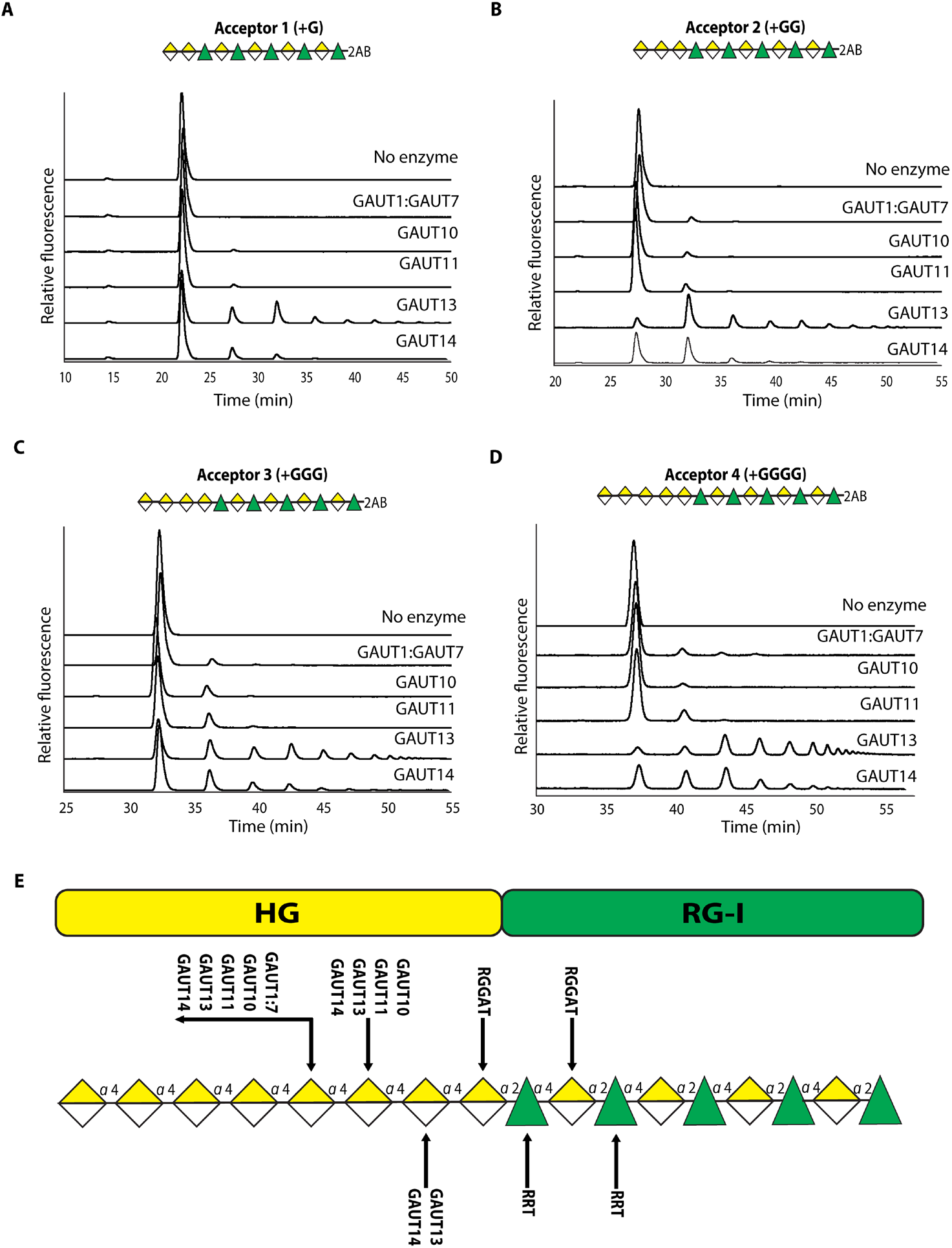
GAUT10, GAUT11, GAUT13, GAUT14, and the GAUT1:GAUT7 complex extend and elongate the HG domain initiated by GAUT13 and GAUT14 in RG-I:HG heteroglycans. (A-D) The different GAUTs were tested for their ability to elongate the RG-I:HG products RG-I(G)-DP8 +G, +GG, +GGG, and +GGGG initiated by GAUT13 and GAUT14 and produced and purified from large-scale enzyme reactions (Figure S2C and D). The acceptor substrates were incubated with GAUT1:7, GAUT10, GAUT11, GAUT13, and GAUT14 in the presence of 100 µM UDP-GalA donor, 10 µM acceptor, 50 mM HEPES at pH 7.2, 0.25 mM MnCl2 for 1 hour. HG domain extension was monitored by anion-exchange chromatography with fluorescence detection of 2-AB-tagged products. GAUT1:7 was able to elongate +GG, +GGG, and +GGGG acceptors, with elongation efficiency increasing with longer acceptor length. In contrast, GAUT10 and GAUT11 elongated all tested acceptors with a preference for +GGG. Both GAUT13 and GAUT14 efficiently elongated all substrates into a series of products, preferentially converting RG-I-DP8 +GGGG. (E) Proposed model for RG-I:HG Biosynthesis. The model depicts the coordinated enzymatic synthesis of the RG-I:HG domain. RRTs and RGGATs sequentially polymerize RG-I by adding rhamnose and galacturonic acid residues onto a growing RG-I polysaccharide backbone (19). GAUT13 and GAUT14 then initiate HG synthesis on the RG-I by adding the first GalA to RG-I(G), followed by elongation of the initiated HG by GAUT10, GAUT11, GAUT13, and GAUT14. The extended HG is then further elongated by GAUT1:7, GAUT10, GAUT11, GAUT13, and GAUT14.

The above results demonstrated that the GAUT1:7 complex, GAUT10, and GAUT11 lack the ability to initiate HG synthesis on RG-I acceptors but can elongate HG on RG-I:HG heteroglycan once it is initiated by GAUT13 or GAUT14. GAUTs 13 and 14 both initiated and elongated HG on RG-I, with the initiation phase occurring more slowly than the elongation phase, consistent with the two-phase model of the pectic polysaccharide elongation (*21*). The varying efficiencies of the different GAUTs to initiate and/or elongate HG in RG-I:HG heteroglycans support the RG-I:HG heteroglycan synthesis model shown in Fig. 3E. In this model, GAUT13 and GAUT14 initiate RG-I:HG heteroglycan synthesis by adding the first α-1,4-linked GalA onto the non-reducing end of RG-I acceptors. GAUTs 10, 11, 13, and 14 then elongate the initiating GalA, with the GAUT1:GAUT7 complex contributing to RG-I:HG heteroglycan elongation once the heteroglycan contains HG of at least two GalA residues in length.

### A motif within the GAUT14 HV1 region is responsible for RG-I:HG extension activity

Among the biochemically confirmed GAUT HG biosynthetic enzymes, only Arabidopsis GAUT13 and GAUT14 initiate HG synthesis on RG-I acceptors, suggesting that GAUT13 and GAUT14 may possess one or more unique structural features enabling RG-I binding and RG-I:HG extension activity. Like the other GAUTs, GAUT13 and GAUT14 are metal-dependent enzymes that catalyze glycosyl transfer through a retaining mechanism (*15, 21, 25*). Structurally, they have a predicted GT-A fold consisting of a single Rossmann fold with alternating β-sheets and α-helices (*21, 25–27*) (Fig. S4 A-B). They possess a predicted conserved GT domain (residues 208-519 and 207-518, respectively), DxD (Asp-X-Asp) donor-binding motif (residues 351-353 and 350-352, respectively), C-terminal histidine residues (His495 and His494, respectively), and three hypervariable regions (HV 1-3). In GT-A fold proteins, hypervariable regions are predicted to contribute to acceptor recognition and substrate positioning within the binding site (*27*).

To identify structural features that contribute to RG-I recognition and RG-I extension activity, we modeled GAUT14 in complex with its substrates using the AlphaFold3 protein-glycan modeling (Model 1) with Chemical Component Dictionary building blocks joined by bondedAtomPairs syntax, an approach validated against stereochemically defined glycan-protein complexes (*28*). To generate catalytically relevant donor-acceptor configurations, an explicit distance restraint was applied between the acceptor GalA and the donor substrate. Because the oligomeric state of GAUT14 could influence substrate binding and affect the modeling, we determined the oligomerization state and molecular weight of GAUT14 using SEC-MALS (Fig. S5A) and AlphaFold-multimer (Fig. S5B-E) (*29, 30*), taking into account that GAUT14 is glycosylated at two of the three sequons predicted by NetNGlyc (N305, N444, N519) (*31*) (Fig. S6A-D). Based on these analyses, GAUT14 was shown to form a dimer (Fig. S5A-E) and was thus modeled as a dimer bound to UDP-GalA donor substrate, Mn^2+^ metal cofactor, and the different acceptors (RG-I-DP5, HG-DP5, UDP-GalA). The resulting model was compared with a different model generated using an AlphaFold2-based molecular docking and molecular dynamics (MD) simulation approach (Model 2), a workflow benchmarked here against the crystal structure of the GT8 enzyme LgtC (*32*) (Fig. S7).

The AlphaFold3 protein-glycan modeling (Model 1) showed Mn^2+^ coordinated by the GAUT14 DxD motif (ASP350 and ASP352), with the divalent metal interacting with the phosphate group of the donor sugar anchoring it within the active site (Fig. 4A-G, Fig. S8A). The model predicted that the three acceptors bind within a shared acceptor-binding pocket composed predominantly of positively charged and polar residues, with their non-reducing ends oriented towards the donor substrate (Fig. 4D-G). The O4 of the GalA in all three acceptors (red spheres on stick) faced toward the donor anomeric carbon (yellow spheres on stick) of the UDP-GalA at catalytically reasonable distances (2.1-2.4 Å) (Black dashed lines) while also remaining in proximity to the donor β-phosphate oxygen (3.6-3.8 Å) (Cyan dashed lines) (Fig. 4E-G). This orientation closely resembles the donor-acceptor geometry observed in some retaining glycosyltransferases (GTs) that operate through an S_N_i-like mechanism (*25*). This donor-acceptor orientation was supported by the AlphaFold2-based molecular docking and molecular dynamics (MD) simulation approach (Model 2) (Fig. S8B-J). Competitive inhibition experiments further supported the presence of a shared binding site for RG-I and HG (Fig. 4H-I), with HG binding more strongly than RG-I (Fig. S9).

**Fig. 4.**
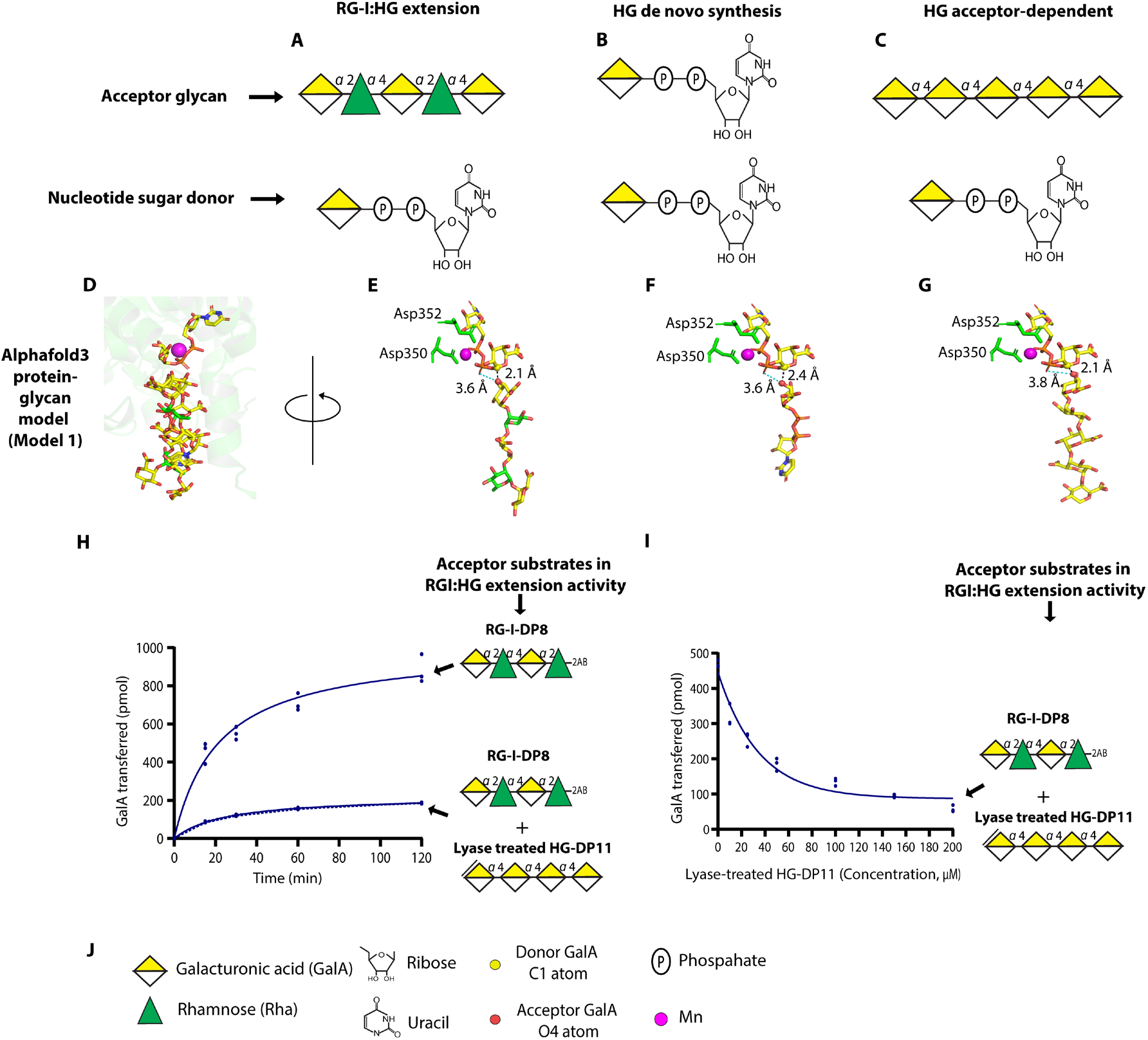
Molecular modeling and competitive inhibition analyses support a shared acceptor-binding pocket for HG, RG-I, and UDP-GalA in GAUT14. (A-C) Schematic representations of acceptor substrates (RG-I-DP5, UDP-GalA, and HG-DP5) and the UDP-GalA donor used for the modeling. (D-G) AlphaFold3 protein-glycan modeling approach was used to predict the binding of Mn^2+^, donors, and acceptors to the active site of dimeric GAUT14. In this model, Mn^2+^ (magenta sphere) was coordinated by the conserved DxD motif (Asp350xAsp352; shown in green), which interacted with the donor UDP-GalA phosphates to anchor the nucleotide sugar in the active site. The three acceptors (RG-I-DP5, HG-DP5, and UDP-GalA) were predicted to bind in the same acceptor-binding pocket, with their non-reducing ends oriented toward the donor substrate. The non-reducing end GalA O4 atom (red sphere) of each of the three acceptors was positioned in proximity to the donor anomeric C1 (yellow sphere) within a catalytic reasonable distance (2.1-2.4 Å; black dashed lines) and near the donor β-phosphate oxygen (3.6-3.8 Å; cyan dashed lines) (E-G), consistent with a retaining GT-A fold enzyme showing an SNi-like mechanism (*25*). (H) Time-dependent competitive inhibition experiment with RG-I:HG extension activity measured in the absence (solid line) and presence (dashed line) of lyase-treated HG-DP11, an inactive HG acceptor that binds but cannot be elongated by GAUT14. The presence of lyase-treated HG significantly reduced RG-I:HG extension activity, suggesting competition for the same binding site. (I) Inhibition assay with 0-200 μM lyase-treated HG showed progressive reduction of RG-I:HG extension activity, confirming that RG-I and HG compete for the same binding site. (J) Symbols used in panels A-I.

Both GAUT14 models predicted that all three acceptors bind in the same acceptor-binding pocket near Hypervariable Region 1 (HV1) (Fig. 5A and Fig. S10). This prompted us to examine the GAUT13 and GAUT14 HV1 region more closely for possible features absent in the other GAUTs. We structurally aligned AlphaFold-predicted GAUT14 (a trifunctional enzyme) with GAUT10 (a monofunctional enzyme lacking RG-I:HG extension activity but having HG acceptor-dependent activity). The GT domains of both proteins aligned well except for their HV1 regions (Fig. 5B and 5C). The alignment showed that the GAUT14 HV1 region had an extended *RTFASKLQSRS* motif (forming an α-helix) that was absent in GAUT10 (Fig. 5B and 5C). This extended region was also present in GAUT13 but absent in GAUT11 (another enzyme lacking RG-I-HG extension activity) and was shortened in GAUT1 (Fig. S11 A-D). To test if the *RTFASKLQSRS* motif mediated RG-I:HG extension activity, we engineered two reciprocal mutant constructs: GAUT14 lacking the motif and GAUT10 with an inserted copy of the motif (Fig. 5D). Recombinant proteins were expressed and purified (Fig. 5E), and enzyme activity measured. As expected, the GAUT14 wild-type enzyme catalyzed RG-I:HG extension, HG acceptor-dependent activity, and HG *de novo* synthesis, while GAUT10 wild-type lacked RG-I:HG extension and HG *de novo* activity (Fig. 5F-I). Remarkably, insertion of the *RTFASKLQSRS* motif into GAUT10 conferred RG-I:HG extension activity to GAUT10 while preserving its native HG acceptor-dependent activity, though it still lacked HG *de novo* synthesis activity (Fig. 5F-I). In contrast, truncation of the same motif from GAUT14 (GAUT14 mut1) eliminated all three enzymatic activities (Fig. 5F-I).

**Fig. 5.**
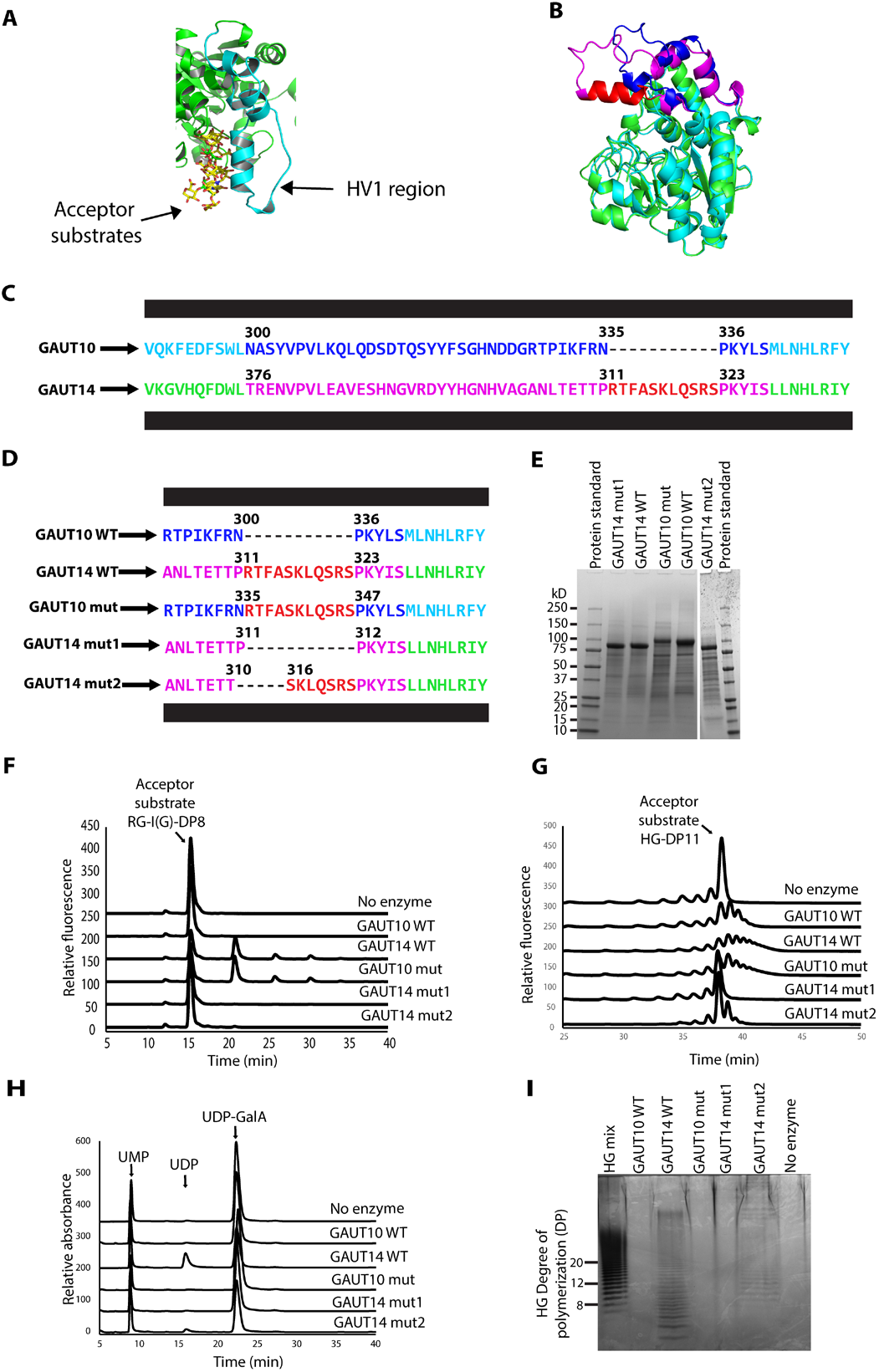
The GAUT14 Hypervariable 1 (HV1) motif *RTFASKLQSRS* mediates RG-I recognition, RG-I:HG extension activity, and is sufficient to provide GAUT10 with RG-I:HG extension activity. Structural comparisons, mutagenesis, and biochemical assays of the GAUT14 HV1 region and its *RTFASKLQSRS* motif reveal the requirement of the HV1 motif for RG-I:HG extension activity. (A) AlphaFold3 protein-glycan model (Model 1) (A) of GAUT14. The model highlights the position of the HV1 region relative to bound acceptor substrates. (B) Structural alignment of GAUT10 (cyan) and GAUT14 (green) GT domains. The GAUT10 HV1 region is highlighted in blue, while that of GAUT14 is in magenta. GAUT14 forms a predicted α-helix (shown in red) within its HV1 region, which is not present in GAUT10. (C) Structure-based sequence alignment of the HV1 regions of GAUT10 and GAUT14, showing the GAUT14-specific “*RTFASKLQSRS”* motif corresponding to the predicted α-helix absent from GAUT10 in panel B. (D) A schematic representation of wild-type (WT) and mutant (mut) constructs of GAUT10 and GAUT14. The GAUT14 mutant (GAUT14 mut1) contains a truncated *RTFASKLQSRS* motif, while the GAUT14 mut2 contains a truncated *PRTFA* motif. The “*RTFASKLQSRS*” motif was inserted into the GAUT10 HV1 region between residues IKFRN and PKYLS. (E) SDS-PAGE analysis of the expressed recombinant proteins (reducing gel). The gel confirms the expression and expected molecular weight of GAUT14 WT (87.9 kDa), GAUT10 WT (91.2 kDa), GAUT14 mut 1(86.9 kDa), GAUT14 mut 2 (87.6 kDa), and GAUT10 mut (92.8 kDa) proteins. (F). Fluorescence chromatograms of RG-I:HG extension activity. GAUT14 WT (1 µM), GAUT14 mut (1 µM), GAUT10 WT (10 µM), and GAUT10 mut (10 µM) were incubated with 100 µM 2-AB-labeled RG-I(G)-DP8 and 1mM UDP-GalA each, with 50 mM HEPES at pH 7.2 and 0.25 mM MnCl2. As expected, the GAUT14 WT protein catalyzed RG-I:HG extension activity, producing a series of products. Also, as previously shown (Figure 2A-B), GAUT10 WT lacked this activity. Interestingly, the GAUT10 mutant with the inserted RTFASKLQSRS motif successfully catalyzed RG-I:HG extension activity, confirming the motif’s role in RG-I:HG extension. (G) Fluorescence chromatograms of HG acceptor-dependent products in a reaction containing 100 µM 2-AB-labeled HG-DP11, 1 mM UDP-GalA, 50 mM HEPES at pH 7.2, 0.25 mM MnCl2. GAUT10 mutants maintain their HG acceptor-dependent activity comparable to the wild-type enzyme. (H) UV detection chromatogram (262 nm) of HG de novo synthesis activity. The chromatogram shows the consumption of 1 mM UDP-GalA (22-24 min) and the formation of UDP (17-19 min), indicating de novo synthesis activity by GAUT14 WT. GAUT10 WT and GAUT10 mutant lack this activity. (I) HP-PAGE analysis of HG de novo synthesis activity. Alcian Blue-Silver staining shows the formation of low-and high-DP HG products in reactions containing GAUT14 WT. No reaction products were observed for GAUT10 WT, GAUT10 mutant, and GAUT14 mut1. This is consistent with what was observed by the UV detection chromatogram in (H) and it suggests that the RTFASKLQSRS motif is specifically involved in RG-I:HG extension, not HG de novo synthesis. The truncation of the RTFASKLQSRS motif in the GAUT14 mutant eliminates all three activities (RG-I:HG extension, HG acceptor-dependent, and de novo synthesis). In contrast, deletion of the four N-terminal residues of the RTFASKLQSRS motif (RTFA) with the preceding Proline residue (P) located at the beginning of the predicted HV1 α-helix was identified as critical and selective for GAUT14 RG-I:HG extension activity since their deletion eliminated only RG-I:HG extension (D-I).

The gain-of-function observed with the GAUT10 mutant suggested that the addition of the HV1 motif (which forms an α-helix) extended the acceptor contact surface, enabling the binding of the less negatively charged RG-I and conferring RG-I:HG extension activity to GAUT10 (Fig. S11E). In contrast, truncating the same region from GAUT14 may have forced the formation of a new junction between amino acid residues that are not naturally positioned adjacent to one another, potentially introducing unfavorable φ/ψ angle constraints that affect the overall protein fold structure of the mutant protein (Fig. S11F). Removing such a large portion of the HV1 α-helix may have disrupted overall enzyme structure rather than specifically affecting RG-I recognition. To test this, we generated a second GAUT14 mutant (GAUT14 mut2) in which the four N-terminal residues of the RTFASKLQSRS motif (RTFA), together with the preceding Proline residue (P), which lies at the beginning of the predicted HV1 α-helix, were deleted (Fig. 5D-I). Unlike the GAUT14 mut1 mutant protein, the GAUT14 mut2 mutant protein was selectively knocked out in GAUT14 RG-I:HG extension activity while both the GAUT14 HG acceptor-dependent and HG *de novo* synthesis activities were preserved (Fig. 5F-I).

Having identified the HV1 motif as a determinant of RG-I extension activity and the shared acceptor-binding pocket for the three GAUT14 substrates, we next sought to identify critical amino acid residues involved in donor and acceptor binding and GAUT14’s three enzymatic activities. Molecular dynamics (MD) simulations of the AlphaFold3 protein-glycan model (Model 1) and the docking/MD-generated model (Model 2) were analyzed using MM/GBSA binding free-energy and hydrogen-bond occupancy calculations to identify candidate residues (*33*) (Table S3A-F and Table S4A-B). Mutating the predicted donor-binding residues D350 and D352 (DxD motif) completely knocked out all three activities in GAUT14, confirming their essential role in binding and anchoring of the donor substrate UDP-GalA (Fig. S12A-H and Fig. S13A-F). Mutations Y478A and K388A eliminated all three activities, suggesting these residues are critical for acceptor binding (Fig. S12A-H and Fig. S13A-F); a conclusion supported by the high score both enzymes had for contacts with all three acceptors (Table S3A-F and Table S4A-B). The possibility that loss of activity resulted from impaired protein folding, however, cannot be excluded. Q498A, Y393A, T310A, N392A, S316A, K449A, Q319A, and R312A mutations affected all three activities, leading to one of several scenarios: a reduction in all three activities, no effect on one of the activities (as in Q319A HG acceptor-dependent activity), or a slight increase in RG-I:HG extension activity (as in S316A and K449A) (Fig. S12A-H and Fig. S13A-F). The results suggest that these residues are part of the binding pocket shared by all three acceptors but they are not essential for enzyme activity. These results place D350 and D352 in the donor site and the HV1 helix along the acceptor groove, where it accommodates all three acceptors, with the PRTFA edge required specifically for RG-I:HG extension (Fig. S14 A-D). Y478 and K388 are additionally required for activity, possibly through their involvement in acceptor binding. The mutants provide targets for testing the biological role of RG-I:HG extension in planta (e.g. the GAUT14 *PRTFA* motif mutant) and for increasing relative RG-I:HG extension via modification of binding site residues (e.g., reducing positive charges in the acceptor binding site, K449A mutant).

### Comparison of GAUT13 and GAUT14 *in vitro* HG *de novo*, HG acceptor-dependent, and RG-I:HG extension activity synthesis rates

Pectins are structurally complex polymers containing multiple pectic glycan domains. GAUT13 and GAUT14, with their three different HG biosynthetic activities, are candidate enzymes for establishing pectin domain structure. To probe the relative contributions of the three different activities to HG and RG-I:HG glycan synthesis *in vitro*, we first established optimal reaction conditions for GAUT14 RG-I:HG extension activity. Our rationale was that since Engle *et al.* (2022) (*15*) had previously shown that GAUT14 has lower HG *de novo* synthetic activity than GAUT13, GAUT14 *de novo* synthesis would compete less for UDP-GalA during catalysis thereby resulting in greater relative RG-I:HG extension activity.

We began by establishing the effects of RG-I acceptor size (RG-I(G) DP 6 to 16 acceptors), pH (6.3 – 7.8), donor substrate concentration (0 – 1000 µM), and acceptor substrate concentration (0 – 200 µM) on GAUT14 RG-I:HG extension activity in 15-minute reactions containing 1 µM GAUT14, which provided linear range data. Previously, Engle *et al*. (2022) (*15*) showed that HG *de novo* synthesis occurs in reactions designed to measure *in vitro* HG acceptor-dependent elongation. We therefore expected that HG *de novo* synthesis was also occurring in the RG-I:HG extension reactions. To account for both activities, we recorded the generation of two products over time. RG-I:HG extension activity was measured as the sum of the peak area of 2AB-labeled extension products, detected by fluorescence (red bars and graphs in Fig. 6A-D). Total activity (RG-I:HG extension activity + *de novo* synthesis activity) was measured as the total amount of UDP produced (indicative of transfer of GalA from UDP-GalA onto the growing product with concomitant release of UDP), detected by UV absorbance (blue bars and graphs in Fig. 6A-D). HG *de novo* synthesis in the same reaction was estimated as the difference between RG-I:HG extension activity and total activity. All activity was reported as pmol GalA transferred.

**Fig. 6.**
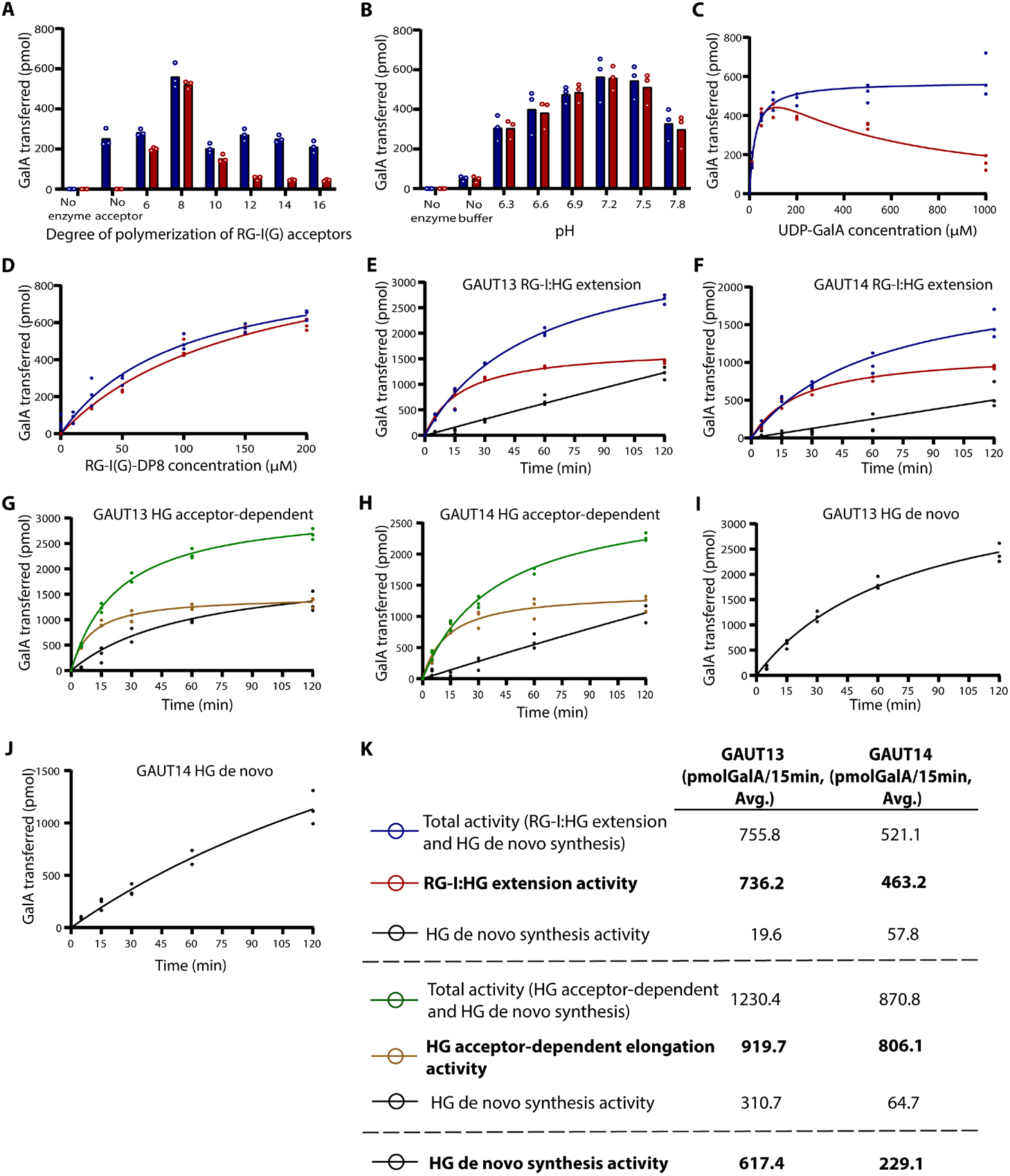
GAUT14 acceptor size preference, pH optimum, and comparison of GAUT13 and GAUT14 RG-I:HG extension activity, HG acceptor-dependent activity and HG *de novo* synthesis activity rates under standard reaction conditions. Standard reaction conditions for GAUT14 RG-I:HG extension activity were established in 15-minute reactions containing 1 µM GAUT14. RG-I:HG extension activity was measured as the amount of 2AB-labelled RG-I:HG extension products detected by anion-exchange chromatography, with separated product peaks quantified by fluorescence detection. Total UDP formed in the same reactions was measured by UV detection and taken as total activity, e.g., RG-I:HG extension + HG *de novo* synthesis activity. Enzyme activity was expressed as pmol GalA transferred. (A) The greatest GAUT14 RG-I:HG extension activity was obtained with RG-I(G)-DP8 acceptor (red bar plot). Total activity (blue bar plot) exceeded RG-I:HG extension activity in all reactions with the difference between these two activities representing HG *de novo* synthesis activity that occurred during the reaction. The smaller difference observed between total and RG-I:HG extension activity in reactions containing the RG-I(G)-DP8 acceptor indicates a greater relative RG-I:HG extension activity compared to *de novo* synthesis activity at 15 minutes with RG-I DP8 as acceptor. (B) The pH optimum for RG-I:HG extension activity using RG-I(G)-DP8 acceptor was pH 7.2. (C) RG-I:HG extension activity increased at low UDP-GalA substrate concentrations (<100 µM) and decreased at concentrations greater than 100 µM. HG *de novo* synthesis in the same reaction was favored at higher UDP-GalA concentrations ≥ 100, plateauing at ∼200 µM. (D) RG-I:HG extension activity increased with increasing concentrations of RG-I(G)-DP8 acceptor. Total activity consistently exceeded RG-I:HG extension activity by only a small amount, suggesting relatively low HG *de novo* synthesis in reactions with RG-I(G)-DP8 acceptor. (E-K) A time-course of the three catalytic activities of GAUT13 and GAUT14 was carried out under standard conditions: 100 µM UDP-GalA (donor), 1 µM enzyme, 50 mM HEPES (pH 7.2), and 0.25 mM MnCl2. RG-I:HG extension activity reactions contained 100 µM RG-I(G)-DP8 acceptor; HG acceptor-dependent activity reactions contained 100 µM HG-DP11; HG *de novo* activity reactions contained only UDP-GalA. For the RG-I:HG extension reactions, total activity (blue), RG-I extension activity (red), and inferred HG *de novo* synthesis activity (black) in the same reaction are shown for GAUT13 (E) and GAUT14 (F). For the HG acceptor-dependent reactions, total activity (green), HG acceptor-dependent activity (brown), and inferred HG *de novo* synthesis activity (black) are shown for GAUT13 (G) and GAUT14 (H). HG *de novo* synthesis activity measured in the absence of an exogenous acceptor (black) is also shown for GAUT13 (I) and GAUT14 (J). For both GAUT13 and GAUT14 at 15 minutes (linear range, K), the greatest activity was observed for HG acceptor-dependent activity, followed by the RG-I:HG extension activity, and then HG *de novo* synthesis, whether inferred from the acceptor-dependent reactions (values not bolded) or measured directly in the absence of an exogenous acceptor (values bolded). As time progressed, for both GAUT13 and GAUT14, the differences between total activity and the RG-I:HG extension or HG acceptor-dependent activities increased (E-H), indicating that HG *de novo* synthesis contributed more to total GAUT13 and GAUT14 catalytic activity at later time points, with GAUT13 showing the highest *de novo* activity. Data represent three independent experimental replicates (n = 3).

The results showed that GAUT14 transfers GalA to all sizes of 2AB-labeled RG-I acceptors tested, with a preference for RG-I(G)-DP8 (Fig. 6A; red bar plot) at a preferred *in vitro* reaction pH of 7.2 (Fig. 6B). The total activity (Fig 6A, blue bar) exceeded the amount of RG-I:HG extension activity, indicating that *de novo* synthesis was occurring simultaneously in all reactions. Curiously, however, apparent *de novo* synthesis activity represented the major activity only in reactions with DP12, 14, and 16 RG-I acceptors. The results suggest that only small RG-I acceptors (RG-I(G)-DP6, 8, and 10) effectively outcompete UDP-GalA and/or *de novo* synthesized HG in the GAUT14 acceptor binding site.

Total activity (i.e., combined RG-I:HG extension and HG *de novo* synthesis activity) increased as UDP-GalA concentration increased, plateauing at ∼200 µM (Fig. 6C). RG-I:HG extension activity in the same reactions increased until 100 µM UDP-GalA, but then decreased from 200 µM through 1000 µM UDP-GalA. These results indicate that HG *de novo* synthesis is favored at higher UDP-GalA concentrations. A possible explanation is that, because the UDP-GalA occupies the same acceptor-binding site as RG-I, as described above (Fig 4D and Fig. S8A-B, F), at high UDP-GalA concentrations, the UDP-GalA outcompetes the RG-I acceptor resulting in less RG-I:HG extension activity product. Another possibility is that accumulation of UDP at high UDP-GalA concentrations preferentially inhibits GAUT14 RG-I:HG extension activity. To test this, RG-I:HG extension reactions were performed in the presence and absence of apyrase, an enzyme that hydrolyzes free UDP. Addition of apyrase significantly increased both RG-I:HG extension activity and total activity in the same reactions, indicating that UDP accumulation inhibits both GAUT14 *de novo* synthesis and RG-I:HG extension activity (Fig. S15). These results show that at UDP-GalA concentrations ≤100 μM, there is a relatively low amount of *de novo* synthesis activity, identifying preferred conditions for testing RG-I:HG extension activity rate *in vitro*.

RG-I:HG extension activity (red) and total activity (blue) both increased with increasing amounts of RG-I(G)-DP8 acceptor (Fig. 6D). Across all RG-I(G)-DP8 concentrations, the total activity only slightly exceeded the RG-I:HG extension activity, indicating that the bulk of the activity was RG-I:HG extension activity when RG-I(G)-DP8 was used as the acceptor.

Using the standard conditions established for GAUT14 RG-I:HG extension activity (100 µM RG-I(G)-DP8 acceptor, 100 µM UDP-GalA (donor), and 50 mM HEPES (pH 7.2)), the catalytic rates of 1 µM GAUT13 and 1 µM GAUT14 for the three different enzymatic activities were compared. Three different types of reactions were carried out: reactions without an exogenous acceptor substrate (for measuring HG *de novo* synthesis); reactions containing 100 µM RG-I(G)-DP8 acceptor (RG-I:HG extension activity assay); and reactions with 100 µM HG-DP11 acceptor (HG acceptor-dependent activity assay). The enzymes were incubated independently under the three reaction conditions, and aliquots of the reactions collected from 0 to 120 minutes were analyzed. In agreement with our prior results, all three enzyme activities (RG-I:HG extension activity, HG acceptor-dependent activity, HG *de novo* synthesis activity) were detected for GAUT13 (Fig. 6E,G,I) and GAUT14 (Fig. 6F,H,J).

During the initial stages of both GAUT13 and GAUT14 *in vitro* synthesis as measured at 15 minutes, HG-acceptor-dependent activity (brown line) (Fig. 6G,H) was the greatest, followed by RG-I:HG extension activity (red line) (Fig.6E,F), and finally HG *de novo* synthesis activity, regardless of whether *de novo* synthesis was inferred from the difference between total and acceptor-dependent activity (blackline) (Fig. 6E-H and 6K) or measured directly in reactions without an exogenous acceptor (Fig. 6I-K). The amount of HG *de novo* synthesis product (black line) continued to increase for at least 120 min. In contrast, the amount of RG-I:HG extension product (red line) (Fig. 6E,F) and HG acceptor-dependent product (brown line) (Fig.6G,H) increased only through 30 minutes and then leveled off. The continued rise in total activity despite the plateauing of exogenous acceptor-dependent activity suggests that HG *de novo* synthesis contributes to progressively more of the total activity over time. The relatively low amount of *de novo* synthesized product produced during the early stages of the reactions indicated that measurement of early initial velocity rates would be the best strategy for measuring the catalytic constants for RG:I:HG extension and HG acceptor-dependent activity so as to reduce the confounding competition for UDP-GalA by the *de novo* synthesis activity. The low HG *de novo* synthesis activity in the early stage of the reaction is consistent with the two-phase distributive elongation model of pectic polysaccharide synthesis proposed for the GAUT1:GAUT7 complex by Amos *et al.* (2018) (*21*). In that model, short-chain oligosaccharides are initially produced during a slow initiation phase of HG synthesis, followed by a more rapid elongation phase once intermediate-length (DP ≥ 11) HG chains are formed. In Fig. 6E-J, the early time points reflect this slow initiation phase, whereas at later time points, the newly synthesized and longer *de novo* synthesized HG chains may compete more effectively for the acceptor-binding site compared to the exogenous RG-I-DP8 and HG-DP11 acceptors.

Taken together, these results show that the three catalytic activities of GAUT13 and GAUT14 contribute to HG product formation at different rates over the course of the reaction. The RG-I:HG extension activity accounts for the majority of the activity in RG-I-containing reactions during the early stages of the reaction for both GAUT13 and GAUT14, but at later stages, the newly synthesized HG (*de novo* synthesized HG) becomes the preferred acceptor over the RG-I acceptors. From a practical framework, these results identified conditions for measuring RG-I:HG extension activity; from a functional standpoint, the results suggest that the initiation of HG *de novo* synthesis proceeds slowly but is followed by a more rapid elongation phase once intermediate-length HG chains are formed. This mechanism is consistent with the two-phase model previously reported for the GAUT1:GAUT7 complex by Amos et al. (2018) (*21*).

### GAUT13/14 trifunctionality is conserved in the seedless non-vascular land plant *Physcomitrium patens*

Having established that Arabidopsis GAUT13 and GAUT14 are trifunctional enzymes that initiate and extend HG on RG-I, elongate HG, and *de novo* synthesize HG, we asked whether this trifunctionality was unique to seed or flowering plants (e.g., Arabidopsis) or whether it was also present in earlier-diverging plants. An NCBI and Phytozome BLAST search using Arabidopsis GAUT13/14 sequences revealed homologs across streptophyte green algae, bryophytes, lycophytes, gymnosperms, and angiosperms (Table S5). We thus tested whether the GAUT13/14 homologs in the non-vascular seedless moss *Physcomitrium patens* also had these three catalytic activities. Of the three GAUT13/14 homologs present in Physcomitrium (Table S5), only two expressed as soluble and active proteins when cloned and heterologously expressed in HEK293F cells (Pp3c23_9440V3 and Pp3c24_12550V3) (Fig. 7A), with Pp3c24_12550V3 yielding less expressed protein as indicated by low protein abundance in the non-reducing gel. Enzyme activity analyses using Pp3c23_9440V3 confirmed its trifunctionality, similar to Arabidopsis GAUT13 and GAUT14 (Fig. 7B-D), with *de novo* synthesis products ranging from low molecular weight HG (HG DP3-DP8) to high molecular weight HG polymers (Fig. 7D) as previously demonstrated for the *de novo* synthesis products of Arabidopsis GAUT1:7, GAUT13 and GAUT14 (*15, 21*). HG acceptor-dependent activity was also detected for Pp3c24_12550V3, although both its HG *de novo* synthesis and RG-I:HG extension activities were relatively low. Taken together, these findings demonstrate that the GAUT13/14 trifunctionality including RG-I:HG extension activity predates the emergence of seed plants and is conserved in early diverging non-vascular land plants.

**Fig. 7.**
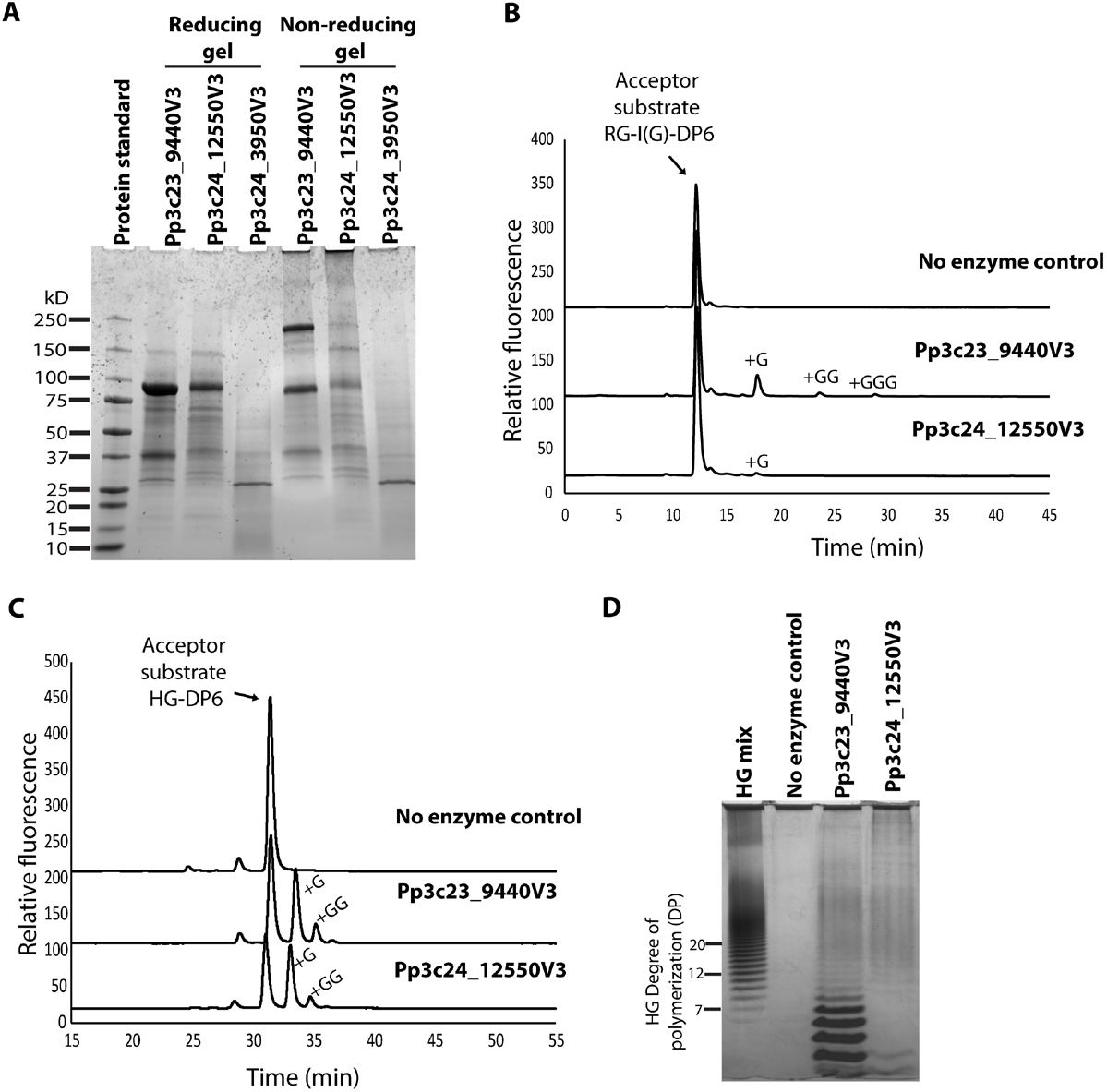
GAUT13/14 trifunctionality is conserved in the seedless non-vascular land plant *Physcomitrium patens*. GAUT13/14 homologs are present in several plant lineages (Table S5), including non-vascular seedless plants such as *Physcomitrium patens*. **(A)** Reducing and non-reducing SDS-PAGE of the three heterologously expressed *Physcomitrium patens* GAUT13/GAUT14 homologs (molecular weights 87.8 kDa for Pp3c23_9440V3 and Pp3c24_3950V3 and 87.7 kDa for Pp3c24_12550V3). Only Pp3c23_9440V3 and Pp3c24_12550V3 yielded soluble and enzymatically active proteins. No band was detected for Pp3c24_3950V3 at the expected monomeric molecular weight. The faint band at ∼87.7 kDa detected for Pp3c24_12550V3 in the non-reducing gel indicated poor expression and low levels of active protein for tests of enzyme activity. **(B)** Fluorescence chromatograms of RG-I:HG extension reaction products separated by anion-exchange chromatography in reactions containing 2-AB-labeled RG-I(G)-DP6 acceptor, 100 µM UDP-GalA donor, 50 mM HEPES (pH 7.2), 0.25 mM MnCl2, 1 µM Pp3c23_9440V3 and 10 µM Pp3c24_12550V3. Pp3c23_9440V3 catalyzed RG-I:HG extension while Pp3c24_12550V3 showed little activity despite being assayed at a high protein concentration. **(C)** Fluorescence chromatograms of HG acceptor-dependent activity using 2-AB–labeled HG-DP6 acceptor confirm that Pp3c23_9440V3 and Pp3c24_12550V3 have HG acceptor dependent elongation activity. **(D)** Alcian blue/silver-stained HP-PAGE of HG *de novo-synthesized* products produced by *Physcomitrium* Pp3c23_9440V3 and Pp3c24_12550V3. Both Pp3c23_9440V3 and Pp3c24_12550V3 produced low-molecular-weight *de novo-synthesized* HG (approximately DP3-DP8 for Pp3c23_9440V3) and high-molecular-weight polymers, as previously documented for GAUT1:GAUT7 (*21*), with Pp3c24_12550V3 having activity than Pp3c23_9440V3. Overall, the trifunctionality observed for Arabidopsis GAUT13 and GAUT14 is conserved in Physcomitrium, with Pp3c23_9440V3 showing higher catalytic activity.

## DISCUSSION

A major challenge in deciphering the function of the plant cell wall is understanding how the different wall polymers are linked to one another in the wall. A key aspect of this is determining how the pectic polysaccharides are linked to form heteroglycans and glycoconjugates in the wall and identifying the enzymes involved in this process. Here, we identified and characterized two trifunctional glycosyltransferases, GAUT13 and GAUT14 as involved in linking the two pectic glycans RG-I and HG. These enzymes were previously known to initiate the production of HG by *de novo* HG synthesis and to elongate HG by acceptor-dependent synthesis. Other catalytic GAUTs such as GAUT1:7, GAUT10 and GAUT11 (including GAUT13 and GAUT14) are able to extend the HG once it is initiated by GAUT13 and GAUT14. However, the other GAUTs lack the ability to initiate HG synthesis on RG-I, i.e., they lack RG-I:HG extension initiation activity. We further showed that GAUT14 preferentially utilizes a DP8 RG-I(G) acceptor substrate at pH 7.2. A confounding factor in carrying out these experiments is that at high UDP-GalA concentrations and long reaction times a significant fraction of the UDP-GalA donor is used for *de novo* HG synthesis. This *de novo* synthesis activity competes for the shared pool of UDP-GalA and negatively impacts the rates of RG-I:HG extension and HG acceptor-dependent elongation. Our structural modeling and experimental results show that HG, RG-I and UDP-GalA acceptor substrates competitively bind to the same binding pocket in GAUT14. Importantly, we show by motif insertion that the GAUT14 hypervariable region 1 (HV1) RTFASKLQSRS motif is sufficient to supply GAUT10 with RG-I:HG extension activity. We further demonstrate that truncation of as little as five residues, PRTFA, from the GAUT14 HV1 region knocks out GAUT14/ RG-I:HG extension activity. These results confirm the essential role of the HV1 region of GAUT13 and GAUT14 in mediating RG-I binding and RG-I:HG extension. The presence of GAUT13/14 homologs in streptophytic green algae and the demonstration of the trifunctional activity of the moss *Physcomitrium patens*, a non-vascular seedless land plant, demonstrate an early evolutionary origin of this trifunctional activity that catalyzes pectic RG-I:HG heteroglycan synthesis, extends HG as a homopolymer, and *de novo* synthesizes HG.

Arabidopsis GAUT13 and GAUT14 share 94% protein sequence identity and are highly expressed in pollen and pollen tubes. Previous studies have shown that they have redundant functions in pollen tubes with *gaut13gaut14* double mutants exhibiting defective pollen tube growth and no recovery of viable plants (*34*). This suggests that GAUT13 and GAUT14 synthesize pectin with critical roles in pollen and pollen tube function. The defective pollen tube phenotype observed in the *gaut13gaut14* double mutants is likely a result of reduced pectin deposition, given that pectins are a predominant component of the cell wall in the tip of growing pollen tubes. RG-I and low-esterified HG regions have been implicated in pollen tube growth (*35, 36*), thus, the RG-I:HG extension activity of GAUT13 and GAUT14 likely contributes to pollen tube growth during fertilization. Whether the bulk of the HG synthesized by these enzymes exists as a homopolymer or is covalently linked to RG-I as part of a heteroglycan remains unknown, since both GAUT13 and GAUT14 can synthesize HG independently and extend RG-I with HG, potentially leading to both polymeric forms of pectin. Since GAUT13/14 catalyzes both homopolymeric HG synthesis and RG-I:HG heteroglycan formation, and because our study shows that HG and RG-I compete for the same binding site, we propose that plants may regulate pectin deposition dynamically through substrate competition in the pollen.

Although GAUT13 and GAUT14 have been shown to have redundant roles in pollen growth, other studies suggest that they may also have unique roles in other stages of plant development. For example, it has been demonstrated that the *gaut13* mutant acts as a suppressor of the pollen tube growth defect in the Arabidopsis alkaline ceramidase *tod1* mutant (*37*). In the *tod1* background, there was an observed increase in turgor pressure in the pollen tube, which may have resulted from excessive pectin deposition in the cell wall. Reducing GAUT13 activity lowered HG synthesis and suppressed the pollen tube growth defects observed in *tod1*. Robichaux et al. (*38*) reported that the *gaut14* mutant suppresses the pollen tube penetration defects of the *Arabidopsis thaliana* O-fucosyltransferase1 (*oft1*) mutant by decreasing pectic HG synthesis. This suppression partially restores silique length, seed set, and pollen transmission in the mutant. The current study shows that the trifunctional activity of Arabidopsis GAUT13/14 is conserved in the early-diverging seedless plant *Physcomitrium patens*. Unlike in Arabidopsis, *Physcomitrium* produces spores rather than pollen, suggesting that this trifunctional activity, which predates the evolution of pollen, has a different activity in spore producing plants. It is likely that the trifunctional activity of GAUT13/14 homologs in *Physcomitrium* represents a distinct developmental or structural role of GAUT13/14 in streptophytic green algae and that this function may also exist in land plants. We hypothesize that only later did a function specific for pollen tube growth in flowering plants emerge.

The results presented here support a model of coordinated pectic HG and RG-I:HG heteroglycan synthesis (Fig. 3E). In this model, RRTs and RGGATs polymerize RG-I by sequentially transferring rhamnosyl and galacturonosyl residues to the non-reducing end of the RG-I backbone (*16, 18, 19, 39*). Once RGGAT incorporates the “final” GalA residue (perhaps based on the length of the RG-I backbone), GAUT13 and GAUT14 recognize the resulting RG-I(G) oligosaccharide ending with a non-reducing-end GalA as an acceptor substrate. These enzymes then add a second GalA, marking the initiation of the HG domain. Previous studies have shown that HG polymer synthesis begins with a slow initiation phase (*21*), and this is applicable to GAUT13 and GAUT14 RG-I:HG initiation activity which initially synthesizes the HG extension on RG-I slowly (Fig. S2 A,B). GAUT10, GAUT11, GAUT13, and GAUT14 contribute by extending the HG in RG-I:HG structure with a third GalA residue, after which GAUT1:7, GAUT10, GAUT11, GAUT13, and GAUT14 continue the elongation of the HG domain. As additional GalA residues are added to the HG domain and the HG reaches an intermediate length, the enzymes transition to a faster elongation phase. This is supported by the series of GalAs added by the GAUTs (especially GAUT13 and GAUT14) when using an RG-I:HG acceptor substrate with at least one added GalA (Fig. 3A-D). The length of the HG domain (GalA units transferred) synthesized by the GAUTs increases until it reaches a size destined for termination, the mechanism of which remains unknown.

The trifunctional activity of Arabidopsis GAUT13 and GAUT14 represents a noteworthy divergence from the other GAUTs, which either exclusively elongate exogenous HG (GAUT10 and GAUT11), both *de novo* synthesize and elongate HG (GAUT1:7), or function as GAUT1 anchors in the Golgi (GAUT5, GAUT6, and GAUT7). This divergence raises two possible evolutionary scenarios for the GAUT evolution. The GAUTs may have gained additional functions over evolutionary time, transitioning from a single-function enzyme (HG acceptor-dependent elongation, an activity present in most GAUTs) to multiple functional enzymes (HG acceptor-dependent elongation, HG *de novo* synthesis, and RG-I:HG extension, as found in GAUT13 and GAUT14). Alternatively, the GAUTs may have evolved via a loss-of-function process in which enzymes like GAUT13 and GAUT14 carrying an ancestral trifunctional activity lost some of those activities over time, such as GAUT1:7 becoming bifunctional (HG *de novo* synthesis and acceptor-dependent elongation) and others such as GAUT10 and GAUT11 becoming monofunctional enzymes (HG acceptor-dependent elongation). Finally, others, such as GAUT5, GAUT6, GAUT7 may have lost all catalytic activity and developed an anchoring function. In either scenario, it is likely that specialization of the biochemical roles of the GAUT family has led to specialization of pectin structure/function. A more detailed structural and phylogenetic analysis of GAUT evolution will be necessary to clarify these potential evolutionary pathways and the functional implications.

An interesting finding from the structural modeling and mutant testing of GAUT14 is the identification of a key motif “*RTFASKLQSRS”* in the HV1 region that is involved in RG-I acceptor binding. The HV1 region, along with HV2 and HV3 present in GTA-fold enzymes, is predicted to be involved in acceptor recognition and specificity (*27*). For example, key residues in the HV2 region of dolichylphosphate mannose synthase 1 (DPM1) create a charged pocket for binding of the phosphate group of the dolichol phosphate acceptor, thereby contributing to the acceptor specificity. Furthermore, the terminal GlcNAc of the N-glycan acceptor is recognized by specific residues in the HV1 region of N-acetylglucosaminyltransferase II (MGAT2) (*27*). The identification of the GAUT14 HV1 *RTFASKLQSRS* motif as both necessary and sufficient to confer RG-I:HG extension activity to GAUT10 provides strong evidence that this motif mediates RG-I recognition. This is supported by the loss of RG-I:HG extension activity when a subset of this HV1 motif *(PRTFA)* is truncated from GAUT14. The gain of RG-I:HG extension activity in GAUT10 upon insertion of the motif highlights how variations in HV1 length and composition can reprogram acceptor specificity within the GAUT family. This suggests that the HV1 region of the GAUTs has roles in acceptor binding, coordination, and specificity. As shown here, mutational analysis can identify amino acid residues whose mutations can be leveraged to provide mutants for probing the biological function of RG-I:HG extension. These mutants are valuable resources for asking questions such as: what is the effect of reduced pectic RG-I:HG heteroglycan on cell wall mechanics, structure, composition, and adhesion? The results reported here provide strong evidence that the HV1 motif is part of the mechanism by which GAUTs evolved new acceptor specificities.

To the best of our knowledge, this is the first study to demonstrate gain-of-function in a GT-A fold glycosyltransferase upon insertion of a hypervariable motif. Previous studies have shown altered substrate specificity, improved enzymatic activity, and/or broader substrate promiscuity when full N-and C-terminal domains of certain GT-B fold proteins were swapped. For example, a study swapped the N-and C-terminal domains of two UDP-sugar glycosyltransferases (UGT) (UGT71C1 and UGT71C2). This resulted in increased catalytic activity and changes in substrate specificity of the chimeric enzymes compared to the parent enzymes (*40*). The chimeric proteins were shown to better glycosylate etoposide compared to their individual parental enzymes. A different study swapped the N-terminal acceptor-binding domain of UGT74B1 and the C-terminal donor-binding domain of UGT74C1 to yield a chimeric protein with enhanced substrate promiscuity (improved glycosylation of structurally diverse thiols and polyphenols) and reduced sensitivity to substrate size (*41*). Another study showed a gain-of-function when a ten-amino acid substitution from the GT-B fold enzyme LanGT1 N-terminal acceptor-binding region (iterative glycosyltransferase involved in landomycin A biosynthesis) was engineered into the non-iterative enzyme LndGT1 (a glycosyltransferase involved in landomycin E synthesis) (*42*). The engineered mutant gained the ability to partially mimic LanGT1 function. However, these studies were done in GT-B fold enzymes, where an entire domain was swapped or multiple amino acids substituted. The results reported here show that small motif addition is sufficient to reprogram acceptor substrate recognition and gain-of-function in GAUTs (GTA fold enzymes).

With the identification of GAUT13 and GAUT14 as pectin biosynthetic enzymes with specific protein domains associated with unique glycan and heteroglycan polymer synthesis, this study supports the hypothesis that unique GAUTs synthesize HG glycan domains in distinct cell wall glycans and/or proteoglycans and that these likely have specialized biochemical and developmental functions (*43*). The findings provide a foundation for probing the roles of different pectic polymers in wall structure, in cellular and plant-level functions such as cell-cell adhesion and cell growth, and in cell wall and cell wall material mechanical and physiological properties with applications in agriculture, bioenergy, and biomaterials.

## MATERIALS AND METHODS

### Heterologous expression of the GAUT proteins

Previously cloned constructs of the transmembrane domain-truncated GAUT proteins (GAUT1Δ167:GAUT7Δ43, GAUT10Δ39, GAUT11Δ39, GAUT13Δ61, and GAUT14Δ61) were used (*15, 21*). The truncated proteins were cloned into mammalian pGEn2-DEST vectors as previously described (*24*) (see Fig. S6A-B) and expressed in HEK293F cells by Transient Transfection (*24*). The secreted proteins were harvested by centrifugation, and the clear supernatant filtered using MF-Millipore^TM^ 5.0 µm MCE membrane filters. The filtered proteins were passed through an AKTA FPLC system with a Diba Omnifit^®^ EZ Chromatography Column packed with QIAGEN Ni-NTA Superflow resin. Bound proteins were washed with a wash buffer of 50 mM HEPES at pH 7.8 containing 15 mM imidazole and 300 mM NaCl. Proteins were eluted in buffer containing 300 mM imidazole.

Eluted proteins were desalted with a GE Healthcare pre-packed desalting column into a 50 mM HEPES storage buffer (pH 7.2) with 10 % glycerol. Recovered proteins were concentrated, and their concentration estimated by Nanodrop UV-visible spectroscopy at 280 nm. The expression, size, and purity of the GAUT-expressed proteins were confirmed by SDS-PAGE (*19*).

### Isolation and purification of RG-I(G) oligosaccharide acceptor substrates of differing lengths

Following the protocol of Amos *et al*., (2022)(*19*), RG-I(G) oligosaccharides with GalA at their non-reducing end were isolated from non-adherent Arabidopsis wild-type (Col-0) seed mucilage. Non-adherent seed mucilage is the outer layer of seed mucilage that disperses upon hydration. Arabidopsis seeds (20 g) were placed into ten 50 mL conical tubes filled with 40 mL deionized water and the tubes were gently rotated overnight and centrifuged at 2500 × g for 5 minutes. The supernatant (containing the mucilage) was filtered through a Millipore Miracloth filter cloth, and lyophilized to yield 335 mg of dry mucilage from the initial 20 gm of seeds. To obtain acid-hydrolyzed RG-I oligosaccharides with GalA at the non-reducing end, dried mucilage was brought to a final concentration of 0.1 M HCl and incubated at 90°C for 32 hours. The mixture was neutralized by addition of 0.1 M NaOH. The resulting oligosaccharide mixture was dialyzed against water using a 3500 Da molecular weight cut-off membrane to eliminate impurities. The sample was dialyzed with three to four water changes over 24 hours, lyophilized, the dry mass measured (128 mg), and the final sample resuspended in water at 20 mg/mL.

Following acid hydrolysis, the oligosaccharides were fluorescently labeled at the reducing end based on previously described protocols (*19*). A solution containing 10% acetic acid (v/v), 0.2 M 2-aminobenzamide, and 1 M sodium cyanoborohydride was added to the sample and incubated at 45 °C for 16 hours. The fluorescently labeled sample was then dialyzed against water and lyophilized to dryness (67.5 mg).

The yield of 2-AB-derivatized oligosaccharides was quantified after resuspension in water by measuring UV absorbance (NanoDrop) and a molar absorptivity coefficient for 2-AB at 338 nm of 2500 M⁻¹cm⁻¹. Individually sized 2-AB RG-I oligomers were obtained by separation over a semi-preparative CarboPac^TM^ PA1 column (22 × 250 mm) and purified as described below. The heterogeneous sample was injected into the HPLC system and individual peaks containing homogeneous RG-I oligosaccharides (DP6-DP18) separated by anion exchange chromatography were collected. Each collected oligosaccharide fraction was dialyzed against water using 3500 Da molecular weight cut-off membrane, lyophilized to dryness, and its dry mass measured. The mass yield of each fraction ranged from 3.2 to 6.6 mg. These were resuspended in water. To determine the purity of the collected fractions, 5 nmol of each fraction was separated over an anion exchange CarboPac™ PA1 column (4 × 250 mm) and the molecular mass of each purified 2-AB-labeled RG-I oligosaccharide (DP6-DP18) was confirmed by MALDI-TOF-MS. All final isolated fractions had purities of 90-95%.

### GAUT enzymes assays

GAUT1Δ167:GAUT7Δ43 (1 µM), GAUT10Δ39 (10 µM), GAUT11Δ39 (10 µM), GAUT13Δ61 (1 µM), or GAUT14Δ61 (1 µM) were independently incubated in 30 µl total reaction volumes containing 50 mM HEPES (pH 7.2), 0.25 mM MnCl_2_, 1 mM UDP-GalA donor, and 100 µM RG-I(G)-DP8 acceptor and the designated amount of enzyme unless otherwise indicated to detect RG-I:HG extension activity. Reactions were incubated for 1 hour and 12 hours, after which the reactions were stopped by boiling for 5 minutes. The samples were analyzed by anion exchange chromatography using a DIONEX HPLC system coupled to an RF-2000 fluorescence detector and by MALDI-TOF mass spectrometry (see below).

For *in vitro* HG acceptor elongation assays, the same concentrations of GAUT enzymes as used above for the RG-I-HG extension assay were incubated in 30 µl reactions containing 50 mM HEPES (pH 7.2), 0.25 mM MnCl_2_, 1 mM UDP-GalA donor, and 100 µM HG-DP11 acceptor. Unless otherwise noted, samples were incubated for 1 hour and 12 hours at 30°C. Reactions were stopped by boiling and analyzed by MALDI-TOF mass spectrometry to confirm HG elongation activity.

To qualitatively detect HG *de novo* synthesized products, 30 µl reactions with the same enzyme concentrations and reaction conditions as described above, except with no exogenous acceptor substrate (i.e., with only UDP-GalA donor substrate), were incubated for 1 hour and 12 hours. Following incubation, the reaction contents were transferred to separate tubes, boiled, and analyzed using high-percentage polyacrylamide gel electrophoresis (HP-PAGE; see methods below).

Standard reaction conditions were established for measuring GAUT14 RG-I:HG extension activity and used to compare the trifunctional enzymatic activities of GAUT13 and GAUT14. Reaction products were analyzed using a CarboPac^TM^ PA1 anion-exchange analytical column (4 × 250 mm) attached to a DIONEX HPLC system and equipped with both a Dionex ICS Series PDS UV and an RF-2000 fluorescence detector. Preferred degrees of polymerization (DP) of the RG-I(G) acceptor, the pH optimum, and the effects of increasing donor and acceptor concentration on GAUT14 RG-I:HG extension activity were determined. The base conditions used were 50 mM HEPES and 0.25 mM MnCl_2_ at pH 7.2, with 100 μM UDP-GalA donor, RG-I(G)-DP8 acceptor and 1 μM enzyme. In each experiment, the specific parameters were varied, for example, for degree of polymerization experiments the DP of the RG-I(G) acceptor ranged from DP6-DP16, pH from 6.3-7.8, donor substrate (UDP-GalA) concentration from 0-1000 μM, and acceptor substrate concentration from 0-200 μM. Each reaction was incubated for 15 minutes, and an aliquot of the reaction was injected into the CarboPac™ PA1 anion-exchange analytical column. These conditions were used to compare the three catalytic activities of GAUT13 and GAUT14 over a time-course assay.

To quantify enzymatic activity, the total amount of UDP detected relative to the amount of UDP-GalA used up during the reaction was determined, which represented the total formation of products due to the transfer of GalA from the UDP-GalA donor to the RG-I acceptor (RG-I:HG extension activity) and transfer of GalA from the UDP-GalA donor to HG produced by either HG acceptor-dependent elongation or HG *de novo* synthesis. The total UDP formed, measured by UV detection of the UDP peak in anion exchange chromatograms of the product produced (see below), was taken as a measure of total enzyme activity and converted to “pmol GalA transferred”. We also accounted for UMP, which can arise from enzymatic or chemical hydrolysis/degradation of UDP during the reaction. The fluorescence detector was used to detect 2-AB-labeled products produced in the same reaction, and the amount of product formed was also represented as pmol GalA transferred. The difference between total activity measured by UDP formation and the corresponding activity for formation of the 2-AB-labeled HG or HG-extended RG-I product was used to infer HG *de novo* synthesis activity in the same reaction.

### Isolation of RG-I:HG Extension products synthesized by GAUT13 and GAUT14 for use as acceptor substrates

The RG-I:HG extension products synthesized by GAUT13 and GAUT14 were isolated and purified for use as acceptor substrates for testing whether other GAUTs could extend the HG domain that was initiated on RG-I by GAUT13 and GAUT14. To achieve this, the RG-I:HG products were produced in 20-hour GAUT13 and GAUT14 RG-I:HG extension reactions, each in a total volume of 4000 μL. Reaction conditions were 2 μM enzyme in a 50 mM HEPES, pH 7.2 buffer containing 0.25 mM MnCl_2_, 1 mM UDP-GalA (donor substrate), and 100 μM RG-I(G)-DP8 (acceptor substrate).

The reaction products were injected into a CarboPac^TM^ PA1 anion-exchange analytical column (4 × 250 mm) equipped with a fluorescence detector. Prominent peaks corresponding to the products were collected in 50 mL Falcon tubes, dialyzed, and lyophilized to dryness. The collected peaks included: RG-I(G)-DP8 (acceptor substrate), RG-I(G)-DP8 + G (one added GalA), RG-I(G)-DP8 + GG (two added GalAs), RG-I(G)-DP8 + GGG (three added GalAs), and RG-I(G)-DP8 + GGGG (four added GalAs). The dried products were resuspended in deionized water and the purity confirmed by anion-exchange chromatography analysis using the Dionex HPLC system. Molecular mass of the products was verified by MALDI-TOF MS.

### Analysis of enzymatic products using anion-exchange chromatography

A CarboPac^TM^ PA1 anion-exchange analytical column (4 × 250 mm) coupled to a Dionex HPLC system equipped with an ICS Series PDS UV detector and an RF-2000 fluorescence detector was used to analyze reaction products. Prior to use, the column was equilibrated with 50% mobile phase A (filtered deionized water) and 50% mobile phase B (1 M ammonium formate) for 5 minutes at a rate of 1 mL/minute. After equilibration, a 15-minute blank was run to remove any preexisting contaminants from the column or HPLC tubing. Reaction samples were diluted in deionized water (or 100 mM ammonium hydroxide for HG reactions to remove lactones), brought to a total volume of 1 mL, and injected into the HPLC system for separation. A linear gradient of 5% to 20% mobile phase B over 5 minutes, 20% to 50% over the next 30 minutes (or 40 minutes for HG acceptor-dependent reaction gradients), and finally 50% to 100% over the last 10 minutes was used with a total run time of 45 minutes (55 minutes for HG acceptor-dependent reaction gradients).

UV absorbance at 262 nm was used to monitor consumption of donor substrates (UDP-GalA) and formation of nucleotide by-products (UDP) via an in-line UV detector. Peaks corresponding to the nucleotide sugar donor and its by-product (23-25 and 17-20 minutes, respectively) were identified based on retention times established from known standards. Fluorescently labeled oligosaccharide products were detected using an in-line fluorescence detector set to medium sensitivity with excitation (330 nm) and emission (420 nm) wavelengths optimized for the fluorescent tag (2-aminobenzamide, 2-AB). Both UV and fluorescence peaks were analyzed using Chromeleon 6.80 software to calculate the percentage area of each product peak(s) relative to the starting substrate. Finally, enzyme activity was quantified based on the moles of substrate converted to product, and this was converted to “pmol GalA transferred”.

### Analysis of enzymatic products using MALDI–MS

MALDI-TOF mass spectrometry was used to determine the molecular masses of oligosaccharides. The matrix, 40 mg/ml 2,5-dihydroxybenzoic acid matrix stock solution was prepared in 50% methanol and this was diluted in a 1:1 ratio with deionized water to produce a ready-to-use matrix solution. Reaction samples were diluted in a 1:4 ratio with either water or ammonium hydroxide (for HG-containing samples, to eliminate lactones).

An aliquot (1 µL) of the diluted samples was mixed with the matrix solution (1 µL) and spotted onto a Bruker MSP 96 Ground Steel MALDI Target Plate, pre-treated with Nafion^TM^ 117-containing solution (Sigma-Aldrich). The spotted samples were dried using a hand-held dryer. The mass spectra were analyzed using a MALDI Bruker Microflex LT spectrometer, and data collected in negative ion mode within a mass range of 500-5000 with a laser frequency of 12 Hz and laser power set to 30-60% of maximum. The resulting spectra of the enzymatic products were examined using Bruker Daltonik FlexControl 3.0 software and peaks corresponding to expected reaction products were identified and compared with control samples. The mass spectra were downloaded into an .emf file.

### Analysis of enzymatic products using high-percentage polyacrylamide gel electrophoresis (HP-PAGE)

Some enzymatic reaction products were visualized by high-percentage polyacrylamide gel electrophoresis (HP-PAGE) using gels containing a resolving gel (30% polyacrylamide, 0.38 M Tris buffer (pH 8.8), 0.035% ammonium persulfate (APS), and 0.18% tetramethylethylenediamine (TEMED)) and a stacking gel (6% polyacrylamide, 0.05% APS, 0.125% TEMED, and 0.13 M Tris buffer (pH 6.8)). Aliquots of the reactions (1-5 μg in 10 uL) were mixed with 2 μl of loading buffer containing phenol red (0.01%), 10% glycerol, and 100 mM Tris pH 6.8 and loaded into the gel wells alongside a molecular weight marker (1 μg of HG ladders of HG DP7-23) and HG standards (0.5 μg each).

After loading into the gel wells, the products were separated by electrophoresis in Tris-Glycine running buffer containing 37 mM Tris and 0.29 M glycine. The gel was electrophoresed at a constant current (17.5-25 mA) until the dye front reached the bottom of the gel. After electrophoresis, the gel was immersed in fixative solution (40% methanol, 10% acetic acid) and stained with 0.1% alcian blue in 40% ethanol for 1 hour. Following staining, excess Alcian blue was removed by extensive washing of the gel with water three times over 12-18 hours. The gel was then stained with Silver Reagent Concentrate (Bio-Rad), developed with Silver Stain Developer (Bio-Rad), and the reaction stopped with 5% acetic acid for 10 minutes. All stained-gel images were captured using Bio-Rad Image Lab 5.2.1 imager with visible light.

### Structure determination of 2-AB-labeled RG-I(G)-DP8+G oligomer by NMR

NMR spectra of 2-AB-labeled RG-I(G)-DP8+G oligomer were acquired on a Bruker NEO spectrometer (^1^H, 799.71 MHz) equipped with a 1.7 mm TCI cryoprobe at 298 and 289 K. The sample (∼0.1 mg) was prepared in D_2_O at pH 7.6 (original) and then at pH 2.7 (adjusted with HCl). The anomeric and H-5 signals were broadened at neutral pH (∼7.6) for the two terminal GalA residues, hampering unambiguous determination of the linkage between them. NMR data were then acquired at lower pH (2.7) where the signals sharpened and it was possible to confirm the α(1→4) linkage between the terminal and penultimate GalAs of the RG-I(G)-DP8+G extension product synthesized by GAUT13 and GAUT14. Structure determination was based on analysis of 1D ^1^H and 2D COSY, TOCSY, NOESY, ROESY, HSQC, HSQC-TOCSY and HMBC spectra. The spectra were processed in Topspin 4.0.5 (Bruker) and analyzed using CCPN AnalysisAssign 3.2.0 (*44*). ^1^H and ^13^C chemical shifts were referenced to the respective 4,4-dimethyl-4-silapentanesulfonic acid (DSS) signals at 0.00 ppm.

### PNGase Digest

The N-glycosylation state of GAUT13 and GAUT14 was determined by digesting both enzymes with PNGase F under native and denaturing conditions. PNGase F was expressed and purified as described (*45*). An aliquot (5 µg) of each protein was incubated with 1 µg of PNGase F under native conditions (250 mM HEPES, pH 7.5, at 37°C for 1 hour) and then placed on ice. Equivalent amounts of GAUT proteins were first reduced and denatured using 40 mM DTT and 0.5% SDS, followed by heating at 100°C for 5 minutes (denaturing condition) so that the effectiveness of PNGase F cleavage of both denatured and non-denatured proteins could be evaluated. The denatured samples were pre-diluted with 1% NP-40 to neutralize the SDS before adding PNGase F (1 μg) and buffer (250 mM HEPES, pH 7.5). Loading buffer was added to the samples, and after incubation, the mixtures were loaded onto an SDS-PAGE gel, and electrophoresis was performed as described above. The degree of deglycosylation was evaluated based on mobility shifts in the gels.

### Size-exclusion chromatography with multi-angle light scattering (SEC-MALS)

A highly purified GAUT14 protein was obtained following separation over a Superose 12 10/300 GL gel filtration size exclusion column. This column was connected to an AKTA FPLC system with a UV detector. An aliquot (100 µg) of the purified protein was separated over a Superdex 200 Increase 10/300 gel filtration SEC prepacked column and the oligomeric state of the protein was determined by a combination of multi-angle light scattering, UV, and refractive index detection. Data analysis was handled with the ASTRA 6.0 software package.

### Molecular modeling: structural prediction, molecular docking, and molecular dynamics simulation

The Molecular docking/molecular dynamics simulated model (Model 2) was created using the AlphaFold2 (*29*) predicted GAUT14 structure, followed by molecular docking (*46*) and molecular dynamics simulation (*47*). We started by predicting the dimeric and trimeric structures of GAUT14 using the full-length GAUT14 amino acid sequence obtained from UniProt(*48*). The highest-ranked models of the dimer and trimer for downstream analysis were selected based on their iPTM+PTM scores. The dimeric form of GAUT14 was used for modeling as described in Fig. S5A-E. To simplify modeling and reduce MD simulation time, we truncated the N-terminal regions of the dimer structure and retained only the glycosyltransferase (GT) domain of GAUT14 (residues 207-518).

To model the substrate binding site of the GAUT14 dimer, we performed a sequential blind molecular docking experiment (the entire GT domain surface searched by the docking algorithm without predefining a binding pocket). First, using the Metal Ion-Binding Site Prediction and Docking Server (MIB2; http://combio.life.nctu.edu.tw/MIB2/ (*49*)), we predicted the ion-binding site for Mn²⁺. We then used the PyRx Virtual Screening Tool (with AutoDock Vina) (*46*) to dock the UDP-GalA donor 3D structure (obtained from the ChemSpider database) on the GAUT14 dimer having the Mn²⁺ bound to the two active sites. Subsequently, we separately docked the following acceptors to generate three distinct coordinate PDB files: UDP-GalA to model HG *de novo* synthesis activity, HG-DP5 to model HG acceptor-dependent activity, and RG-I(G)-DP5 to model RG-I:HG extension activity. Shorter acceptor oligosaccharides (DP5) were used to allow for more precise docking prediction since longer oligosaccharides had more degrees of freedom, making docking less reliable due to increased conformational variability (*50, 51*). Both the HG-DP5 and RG-I(G)-DP5 3D structures were designed and obtained from the GLYCAM-Web tool (https://glycam.org/cb/) (*52*). The top-ranked docked complexes with the lowest binding energy were selected for further analysis.

Each of the three docked complexes containing Mn^2+^, UDP-GalA donor substrate, and their respective acceptors, was subjected to molecular dynamics (MD) simulations to refine the docked poses, confirm that the substrates occupied energetically favorable binding sites, and generate a stable static model. The docked complexes were first prepared by adding missing residues and atoms. Using the CHARMM-GUI solution builder tool (*47, 53–55*), the MD simulation systems were prepared through the following steps: solvating the complexes with explicit water (solvated in a cubic box with TIP3P water molecules), neutralizing the system by adding ions such as K^+^ and Cl^-^, and generating input files for energy minimization, equilibration, and MD production steps. We parameterized the protein-carbohydrate complexes using the CHARMM36M force field, and the carbohydrate topologies were all generated using the CHARMM-GUI.

We performed energy minimization, equilibration, and MD production using the installed AMBER software (Amber/22.0-foss-2021b-AmberTools-22.3-CUDA-11.4.1) at the University of Georgia Sapelo2 Cluster-Georgia Advanced Computing Resource Center (GACRC). First, energy minimization was performed to remove steric clashes until energy convergence was achieved (maxcyc set to 25000). The first 2,500 cycles used the steepest descent method (ncyc=2500), followed by the conjugate gradient method for the remaining cycles if needed. After minimization, an equilibration step was performed, followed by a production step (MD simulation) under NPT (constant pressure, temperature, and the number of particles). The production step was conducted for 500 ns until a stable structure was obtained (monitored using root mean square deviation (RMSD)). The non-bonded cutoff was set to 12 Å. Structure figures were generated using PyMOL. Binding free energy calculations were performed using the Molecular Mechanics/Generalized Born Surface Area (MM/GBSA) (*56*) approach on the MD simulation trajectories, and hydrogen bond occupancy was calculated using the VMD tool. The model produced using this approach was referred to as Model 2 in the results section.

The AlphaFold3 protein-glycan model (referred to as Model 1 in the results section) was generated using the technique described by *Huang et al.,* (2025)(*28*).

### Generation of mutants using site-directed mutagenesis

The top candidate residues showing the strongest interactions with UDP-GalA, HG, and/or RG-I acceptors were mutated to alanine using PCR-based site-directed mutagenesis with primers containing the desired substitutions (Table S6). Additionally, the D350 and D352 residues from the conserved donor-binding DxD motif were also mutated. The mutations were introduced into GAUT14 using a GAUT14:pDONR^TM^221 construct, that was originally constructed by Engle *et al*., 2021(*15*). The mutated genes were then transferred into a destination vector (pGEn2-DEST expression vector) via an LR-Clonase II enzyme reaction. Mutation verification was performed by whole-plasmid sequencing, and the resulting proteins were expressed in HEK293F cells and purified using Ni-NTA affinity chromatography. The trifunctional activities of the mutant proteins were tested and compared to the wild-type GAUT14 protein.

To investigate the functional role of the *RTFASKLQSRS* and the *PRTFA* motifs within the HV1 region of GAUT14 proteins, codon-optimized GAUT14 and GAUT10 wild-type and mutant gene sequences were synthesized by Twist Bioscience® and received in the pTwist-ENTR-Kozak vector. The constructs were subsequently transferred into the mammalian expression vector pGEn2-DEST via Gateway LR recombination cloning to enable recombinant protein expression in HEK293F cells. All constructs were confirmed by whole plasmid sequencing prior to expression.

## Acknowledgments

We thank Lubana Shahin for the gift of lyase-treated HG-DP12, Daniel Tehrani for help with the SEC-MALS analysis, Paul J. Glatz for UDP-GalA substrate, and Alex Lee and Eliza Gazaway for assistance with MM/GBSA troubleshooting. We thank Robert Woods, Natarajan Kannan, Ian Wallace, Jim Leebens-Mack, and Kristen Engle for helpful discussions regarding the research and manuscript.

## Author contributions

Conceptualization: C.T.O., R.A., D.M.

Methodology: C.T.O., R.A., D.M., P.J., C.H., J. V.

Investigation: C.T.O., R.A., J. V.

Visualization: C.T.O, C.H., D.C., J.E., J. V.

Supervision: P.A., K.W.M., D.M

Writing—original draft: C.T.O., D.M

Writing—review & editing: C.T.O., R.A., D.M., P.J., C.H., D.C., J.E., J. V., P.A., K.W.M.

## Supplementary Figures

**Fig. S1.**
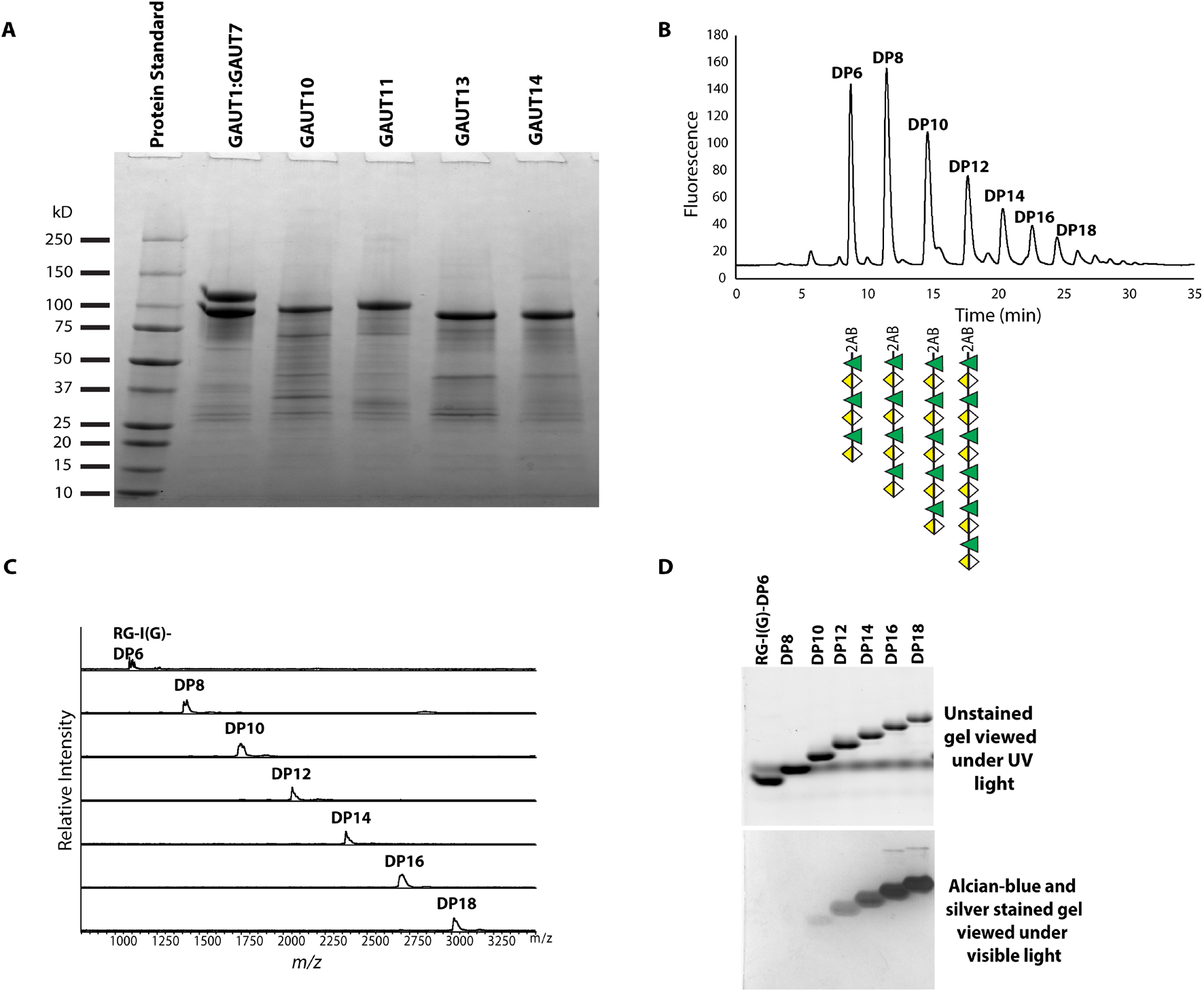
Analysis of expressed GAUT proteins and purification of uniquely sized RG-I oligosaccharide acceptor substrates. **(A)** Expression and molecular weight of Ni^2+^-NTA purified GAUT proteins under reducing conditions (+DTT) confirmed by SDS-PAGE. Major bands were resolved near the expected molecular weights for the GAUT1Δ167 (92.6 kDa):GAUT7Δ43 (99.6 kDa) complex, GAUT10Δ39 (91.2 kDa), GAUT11Δ39 (91.1 kDa), GAUT13Δ61 (88.1 kDa), and GAUT14Δ61 (87.8 kDa). **(B)** Anion-exchange chromatogram with fluorescence detection of purified 2AB-labeled RG-I(G) oligosaccharide acceptor substrates (purified from Arabidopsis seed mucilage) that were used as acceptor substrates for enzyme reactions; acceptors with degrees of polymerization (DP) of 6 to 18 are shown. Glycan symbol representations of RG-I DP 6, 8, 10 and 12 are also shown using Symbol Nomenclature for Glycans (SNFG) (*57*). RG-I(G) is designated as RG-I oligosaccharide with a GalA residue on the non-reducing end. **(C)** MALDI-TOF mass spectra of the purified 2AB-labeled RG-I(G) oligosaccharides (DP6–DP18) illustrated in (B). **(D)** The purity and electrophoretic mobility of the 2-AB-labeled oligosaccharides were analyzed by high-percentage polyacrylamide gel electrophoresis (HP-PAGE). Following electrophoresis, the gel containing size-separated oligosaccharides was first viewed under UV light (top gel) to detect 2-AB-labeled fluorescent oligosaccharides by UV transillumination at 302 nm, then stained with alcian blue and silver reagents and re-imaged under visible light to visualize the stained negatively charged oligosaccharides. RG-I oligosaccharides of ≤ DP8 have a low mass to charge ratio, hence, they do not stain well by alcian blue staining and are not detected by this staining method.

**Fig. S2.**
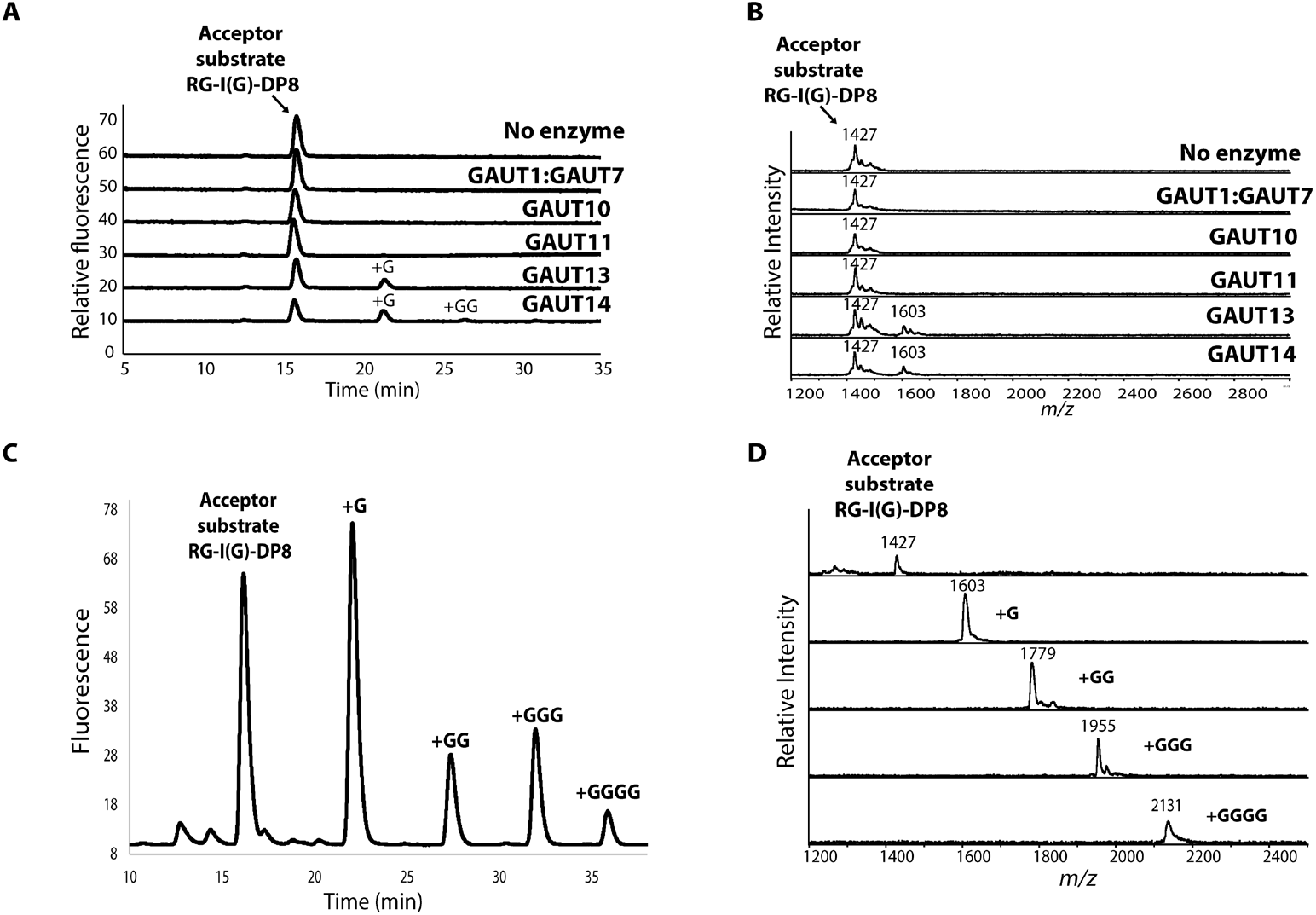
GAUT13 and GAUT14 initiate α(1→4) linked HG synthesis on the non-reducing end of RG-I. **(A)** Anion-exchange chromatogram of GAUT-catalyzed reaction products produced in 1-hour reactions. Reactions contained 50 mM HEPES at pH 7.2, 0.25 mM MnCl2, 100 µM 2AB-labeled RG-I(G)-DP8 acceptor, 1 mM UDP-GalA donor substrate, and 1 µM (GAUT1:7, GAUT13, and GAUT14) or 10 µM (GAUT10 and GAUT11) protein. **(B)** MALDI-TOF MS spectrum of the GAUT-catalyzed 1-hour reaction products shown in (A). **(C)** Separation of large-scale GAUT13 and GAUT14 2-AB-labeled RG-I:HG extension reaction products. The RG-I-DP8 +G, +GG, +GGG, and +GGGG products were resolved and separated by anion-exchange chromatography with fluorescence detection and size-purified. The reaction (20 hours) contained 50 mM HEPES, pH 7.2, 0.25 mM MnCl2, 1 mM UDP-GalA (donor substrate), 100 μM RG-I(G)-DP8 (acceptor substrate), and 2 μM GAUT13/GAUT14. **(D)** MALDI-TOF MS analysis of the purified large-scale GAUT13 and GAUT14 RG-I:HG extension reaction products.

**Fig. S3.**
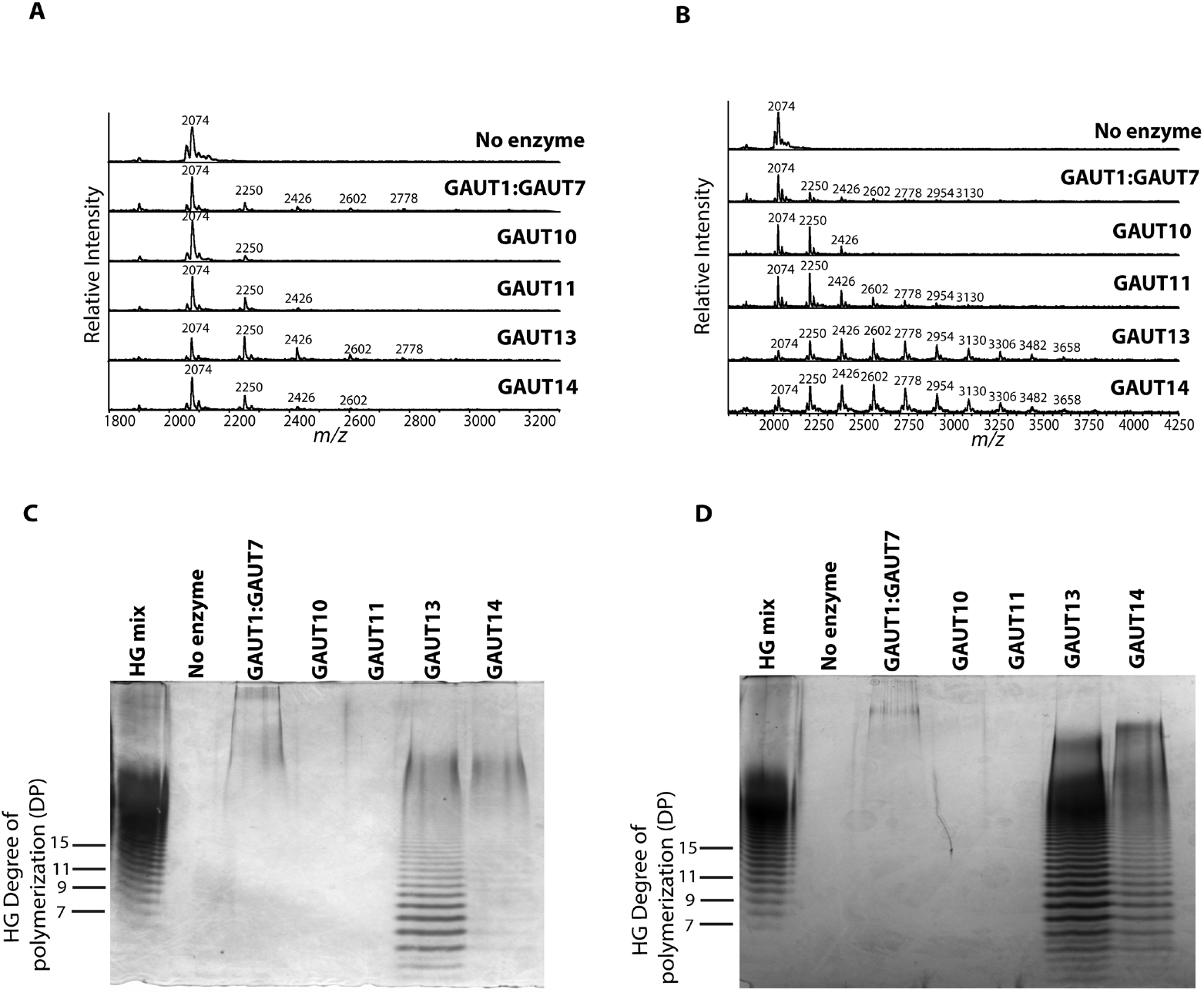
Confirmation that some GAUTs elongate HG oligosaccharides and/or *de novo* synthesize HG *in vitro*. **(A-B)** MALDI-TOF MS analysis of HG elongation products synthesized on HG DP11 acceptor by GAUT1:GAUT7 complex, GAUT10, GAUT11, GAUT13, and GAUT14 after 1 hour (A) and 12 hours (B). The reaction conditions were 50 mM HEPES (pH 7.2), 0.25 mM MnCl2, 1 mM UDP-GalA donor, 100 µM 2AB-labeled HG DP11 acceptor, 1 µM GAUT1:GAUT7, GAUT13, and GAUT14; 10 µM GAUT10 and GAUT11. **(C-D)** Alcian blue and silver-stained high-percentage polyacrylamide gel electrophoresis (HP-PAGE) showing that the GAUT1:GAUT7 complex, GAUT13, and GAUT14 *de novo* synthesize HG in the absence of an exogenous acceptor substrate, after 1 hour (C) and 12 hours (D) of reaction. The gel shows that GAUT1:GAUT7 has HG:GalAT activity, resulting in high-molecular-weight polymeric HG even after 1 hour, whereas GAUT13 and GAUT14 synthesize a series of differently-sized HG oligomers and polymers over 1 and 12 hours, agreeing with the prior results of (*15*). The same reaction conditions as in Figure S3A-B were used in Figure S3C-D, except that no acceptors were included.

**Fig. S4.**
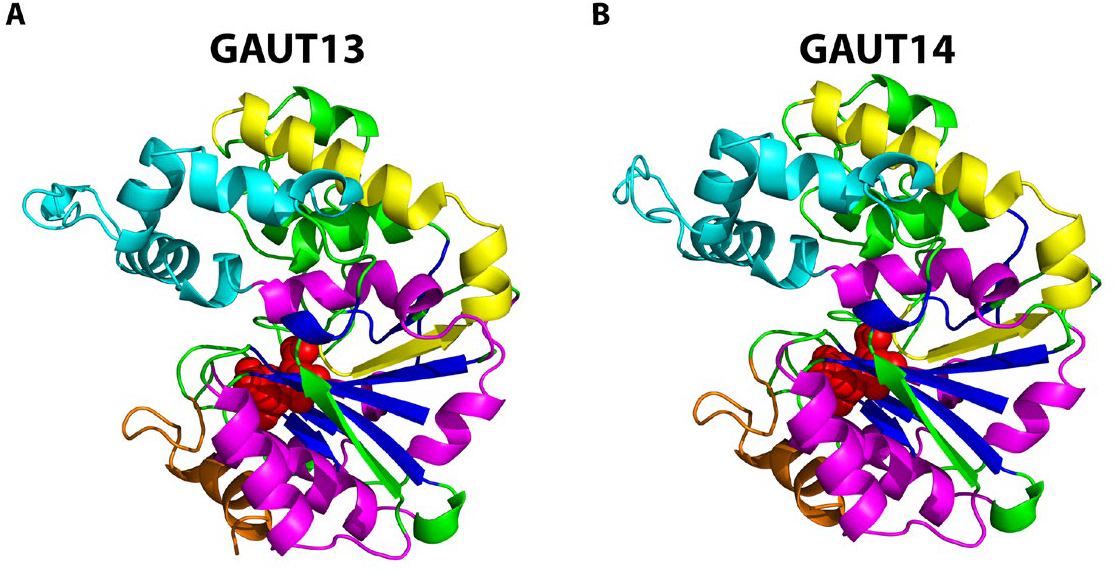
Structural annotation of the AlphaFold3-predicted structures of GAUT13 and GAUT14. **(A-B)** Cartoon representation of the predicted GT-A fold structures of GAUT13 and GAUT14 monomers predicted by AlphaFold3. Each monomer has a single Rossmann fold with eight alternating β-sheets and six α-helices (blue and magenta, respectively) (*27*); https://uga-gta.netlify.app/). The GT domain in GAUT13 and GAUT14 spans amino acid residues 208-519 and 207-518, respectively, and includes a predicted DxD donor-binding motif (residues 351-353 for GAUT13 and 350-352 for GAUT14; red spheres), and the Hypervariable regions (HV1 in cyan cartoon, HV2 in yellow cartoon, and HV3 in orange cartoon).

**Fig. S5.**
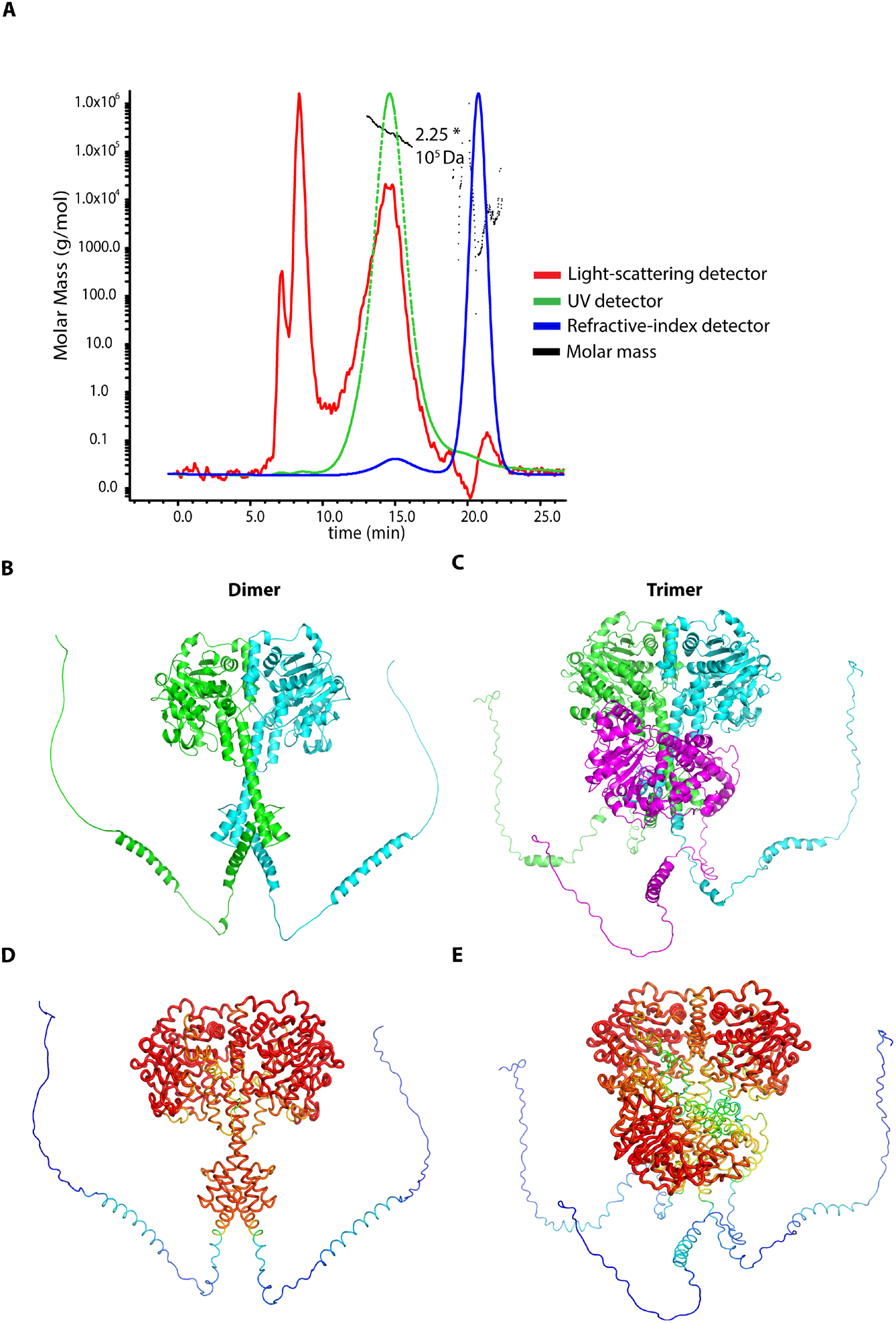
SEC-MALS analysis and AlphaFold structural prediction of the GAUT14 oligomeric state. **(A)** Size-exclusion chromatography coupled with multi-angle light scattering (SEC-MALS) analysis of GFP:GAUT14 revealed a molecular mass of 225 kDa (black dotted line), a molecular weight value between predicted homodimer (176 kDa) and homotrimer (264 kDa). SEC-MALS: light scattering (red line), UV (green line), refractive index (blue line). **(B-E)** AlphaFold Multimer was used to predict the potential dimer and trimer structures of GAUT14. Predicted structures of the GAUT14 homodimer and homotrimer are shown as cartoon representations (top panel), with dimer subunits colored green and cyan (B), and trimer subunits colored green, cyan, and purple (C). In the dimer, the domains form a symmetrical interface, while the trimer appears more asymmetrical (B,C). The confidence level of each predicted subunit and their interacting interface in the GAUT14 dimer complex (by PyMOL’s B-factor putty representation) is high compared to the trimer structure (red indicates high confidence and blue indicates low confidence) (D,E). The results support GAUT14 dimer stoichiometry. The N-terminal region (containing the transmembrane domain and signal peptide) of each subunit in the dimer and trimer structures appears blue because it is not well predicted by AlphaFold3.

**Fig. S6.**
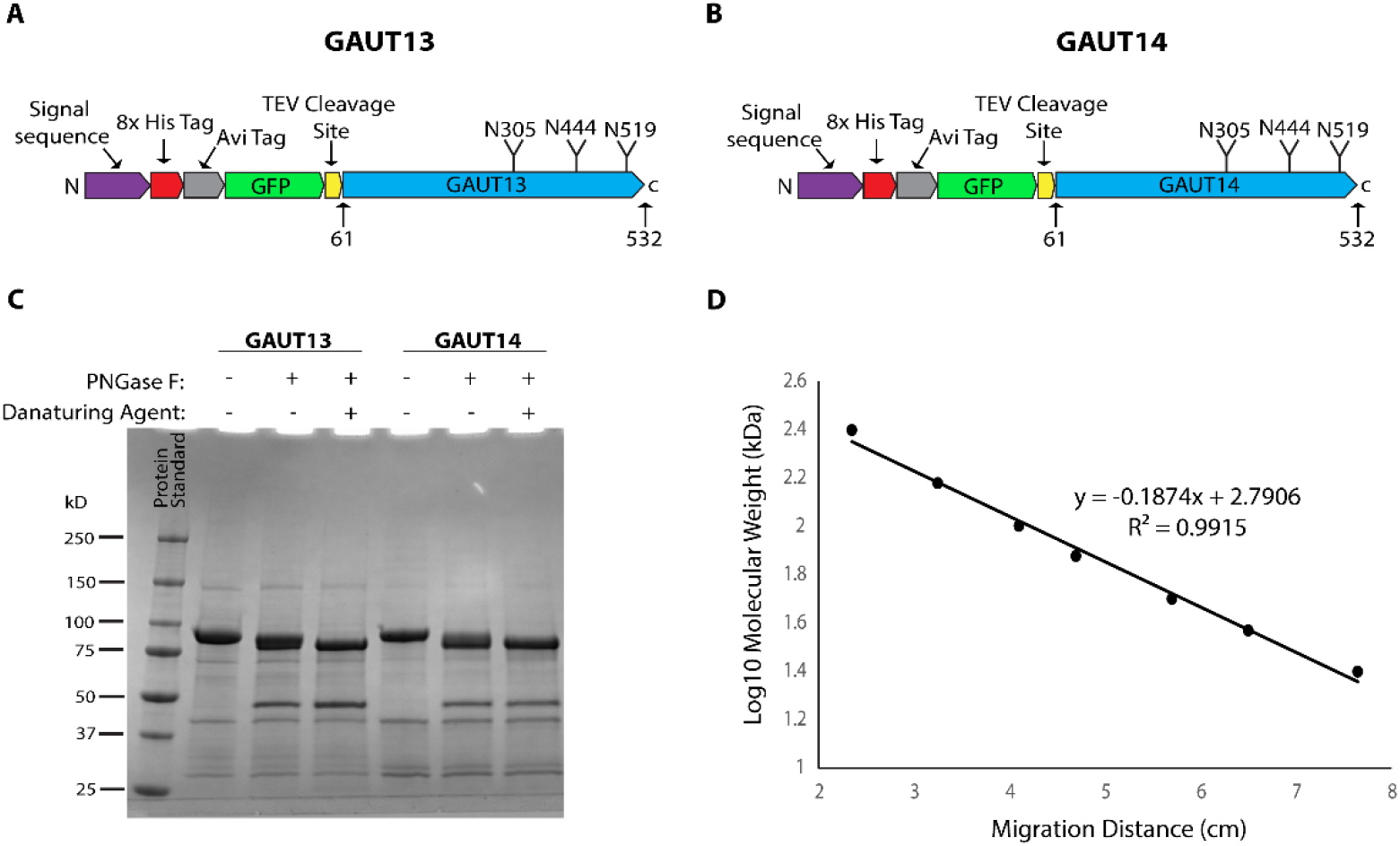
GAUT13 and GAUT14 are lightly N-glycosylated. **(A-B)** Schematic representation of GAUT13 and GAUT14 constructs used in this study, with the location of their predicted N-glycosylation sites using the NetNGlyc N-Glycosylation prediction server (*31*). The constructs in the PGEN2 expression vector contain a signal sequence, 8x His tag, Avi tag, GFP, TEV cleavage site, and the protein sequence. Three N-glycosylation sites (N305, N444, and N519) within the GAUT13 and GAUT14 GT domains were predicted. **(C)** SDS-PAGE reducing gel showing the effect of PNGase F treatment on N-glycosylation of GAUT13 and GAUT14. GAUT13 and GAUT14 were treated with PNGase F under both native and denaturing conditions. Under native conditions, proteins were incubated with PNGase F (+) without denaturing agents (-). Under denaturing conditions, proteins were first treated with DTT and SDS (+), followed by heating at 100°C for 5 minutes, and then treated with PNGase F (+). The slight mass shift in treated vs untreated GAUT13/14 indicates that both proteins are lightly N-glycosylated. **(D)** Semi-log plot of protein standard (ladder). The migration distances (cm) were plotted against the log molecular weights (kDa) of the protein standard in C. The calibration curve was used to calculate the molecular weights of GAUT13 and GAUT14 before and after PNGase F treatment. Using the plot-based regression equation (y = –0.1874x + 2.7906, R^2^ = 0.9915), the untreated protein migrating at 4.5 cm corresponded to ∼88.6 kDa, while the PNGase F-treated protein migrating at 4.6 cm corresponded to ∼84.8 kDa. The ∼3.8 kDa decrease in molecular weight is consistent with the removal of ∼2 N-glycans (average plant N-glycan ∼2 kDa (*58*)), suggesting that GAUT13 and GAUT14 are modified at two of the three predicted N-glycosylation sites.

**Fig. S7.**
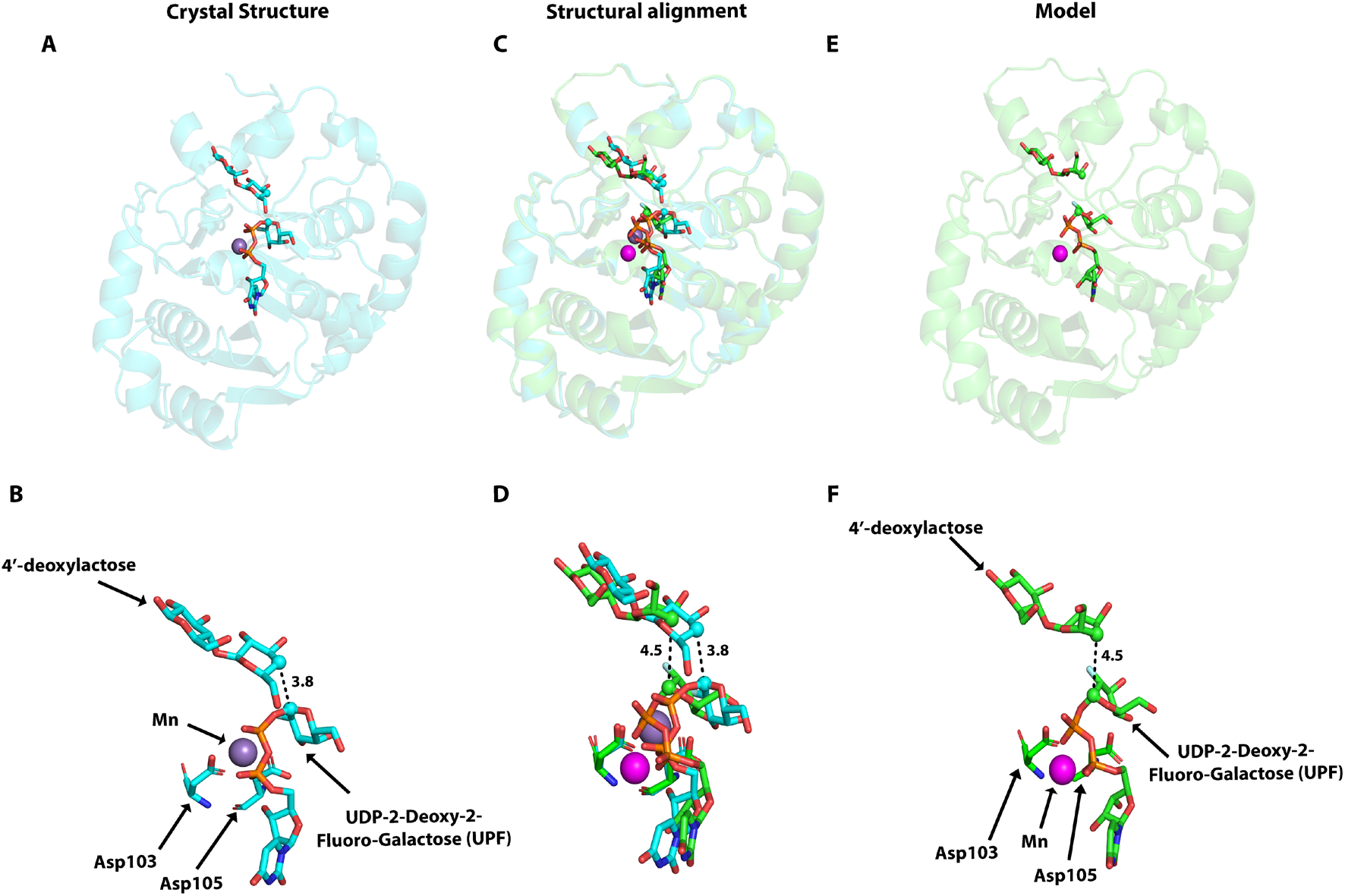
Validation of the molecular docking and molecular dynamics (MD) simulation workflow using the GT8 enzyme LgtC. (A-F) To evaluate the reliability of the molecular docking and molecular dynamics (MD) simulated models (Model 2) used in this study prior to modeling GAUT14, we first modeled LgtC, a well-characterized glycosyltransferase from *Neisseria meningitidis*. LgtC is a retaining GT in the same GT8 family as the GAUTs (*32*). The experimentally determined crystal structure of LgtC in complex with cofactor Mn^2+^, UDP-2-deoxy-2-fluorogalactose (UPF), and the acceptor analog 4′-deoxylactose (PDB ID: 1GA8) is shown in (A-B). The Mn^2+^ ion is coordinated by the DxD motif (Asp103 and Asp105), which interacts with the donor phosphate group. AlphaFold-2-predicted, molecular docked/MD-simulated LgtC model showing Mn^2+^ coordination by Asp103 and Asp105 and positioning of UDP-2-deoxy-2-fluorogalactose and 4′-deoxylactose within the active site (E-F). Comparison of the modeled structure with Mn^2+^, donor, and acceptor sugar analogs in its binding site with the experimentally determined 3D structure of LgtC showed that the experimentally determined and modeled structures and complexes aligned very closely (high root-mean-square deviation (RMSD) of 0.157 Å) (C-D). The black dashed lines indicate distances between the donor anomeric carbon (C1) and the acceptor C4 position. These distances appear relatively long in both the crystal structure and the modeled complex because the acceptor analog 4′-deoxylactose (used to prevent catalytic sugar transfer during crystallization) lacks the C4 hydroxyl group required for glycosyl transfer. Overall, these results support the reliability of the computational approach to model the GAUT14 substrate complex.

**Fig. S8.**
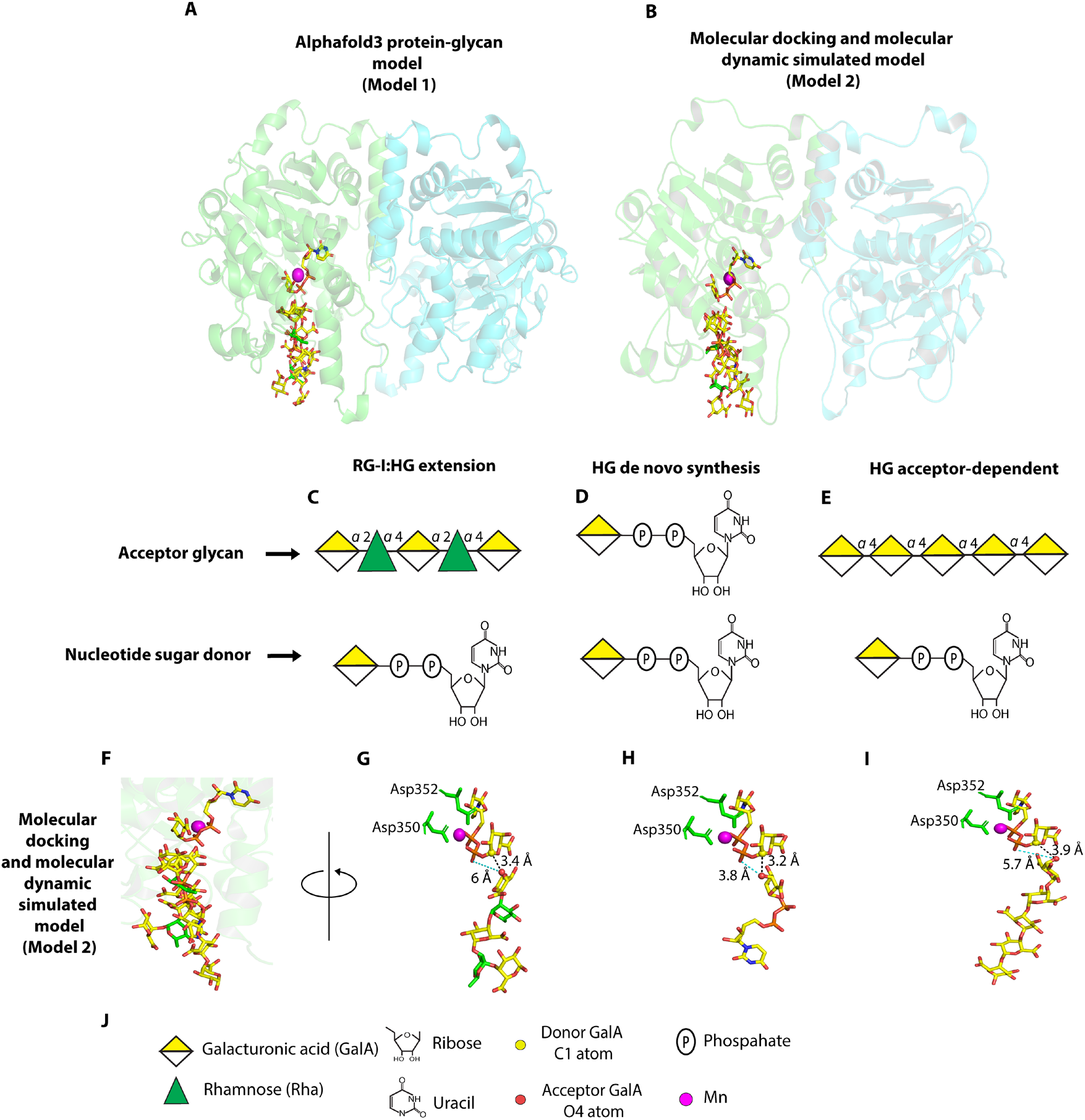
Molecular modeling of GAUT14 donor and acceptor binding site. **(A-B)** AlphaFold3 protein-glycan model (Model 1) and molecular docking-and molecular dynamics-generated model (Model 2) of the GAUT14 GT-Domain dimer with bound substrates. The homodimer chains are represented in green (left) and cyan (right) cartoons. Bound substrates include the UDP-GalA donor substrate, the acceptor substrates RG-I-DP5, HG-DP5, UDP-GalA, and the metal Mn^2+^. **(C-E)** Schematic representations of acceptor substrates (RG-I-DP5, UDP-GalA, and HG-DP5) and the UDP-GalA donor used for the modeling. **(F-I)** The docking/MD model (Model 2) was similar to the AlphaFold3 protein-glycan model (Model 1) (Fig. 4D-G) except that it placed the O4 in the GalA of the three acceptors farther from the donor C1 atom (3.2-3.9 Å) and β-phosphate oxygen (3.8-6.0 Å). Additionally, its HG oligomer (I) had its non-reducing end GalA O4 atom oriented away from the donor in a catalytically unfavorable configuration. **(J)** Symbols used in panels A-I.

**Fig. S9.**
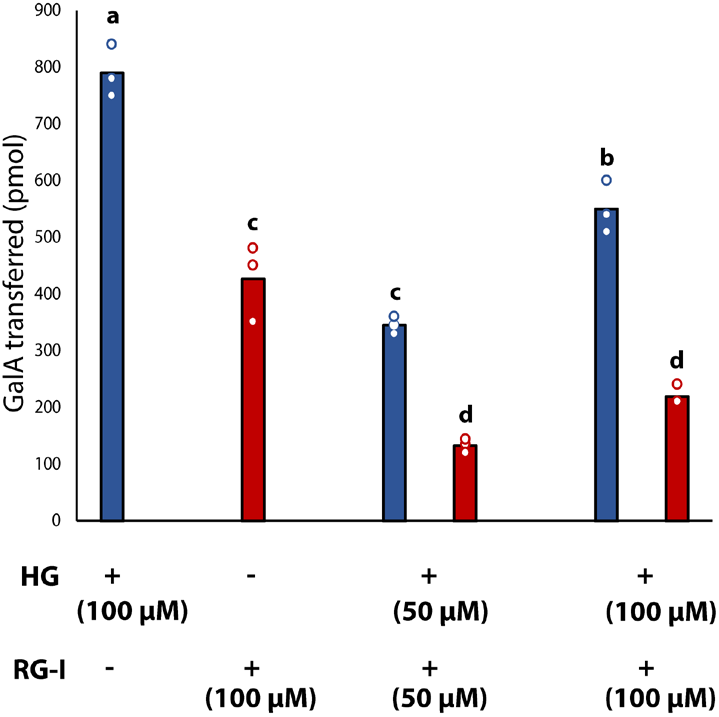
Comparison of GAUT14 activity in the presence of individual or combined RG-I(G)-DP8 and HG-DP11 acceptors. GAUT14 enzyme (1 µM) was incubated independently in solutions containing 100 µM UDP-GalA, 50 mM HEPES at pH 7.2, 0.25 mM MnCl2, 2-AB-labeled HG-DP11 (100 µM or 50 µM) for HG acceptor-dependent activity, and 2-AB-labeled RG-I-DP8 (100 µM or 50 µM) for RG-I:HG extension activity. Reactions with only 100 µM HG-DP11 acceptor had significantly higher GalAT activity than those with only 100 µM RG-I(G)-DP8 acceptor for GAUT14. In the presence of both acceptors, HG acceptor-dependent activity was significantly greater than RG-I:HG extension activity, demonstrating that HG binds more tightly to GAUT14 than RG-I. Reactions were carried out for 15 minutes. Error bars represent the standard deviation of n = 3 independent experiments. Different letters above bars indicate statistically significant differences between conditions (P < 0.05).

**Fig. S10.**
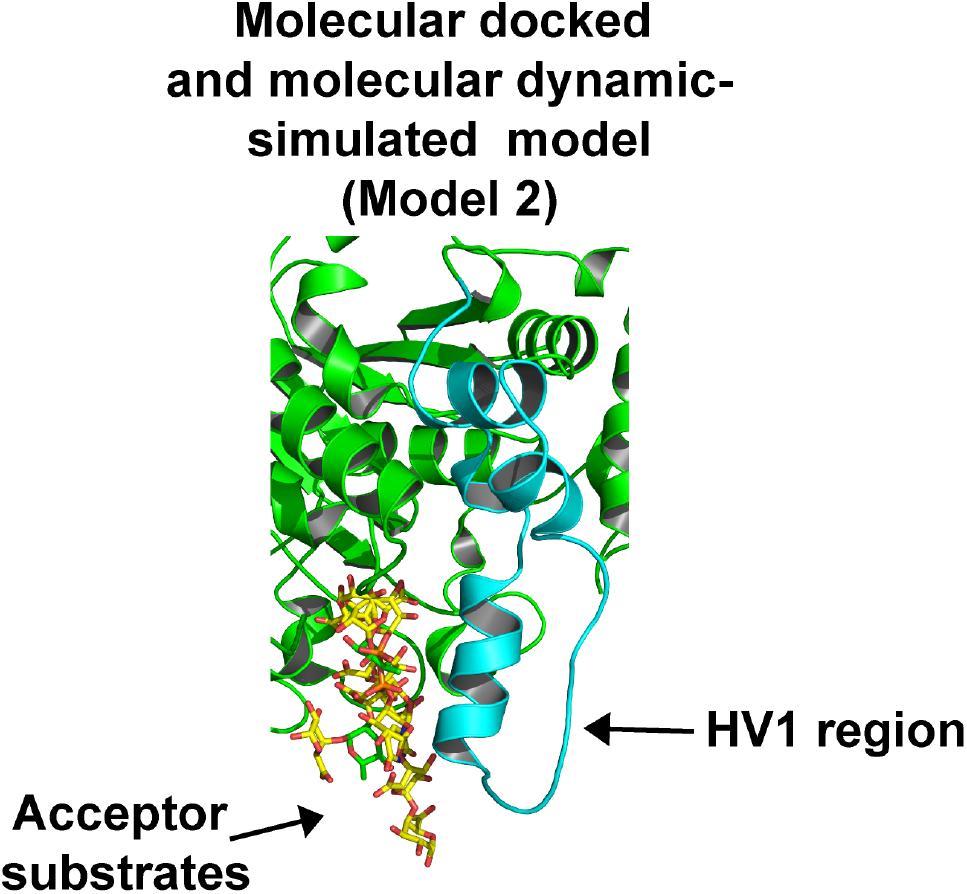
GAUT14 HV1 region position relative to bound acceptor substrates in the molecular docking/MD-simulated model (Model 2). Molecular docking/MD-simulated model (Model 2) of GAUT14 highlighting the position of the HV1 region relative to the bound acceptor substrate, consistent with what was observed in the AlphaFold3 protein-glycan model (Model 1; Fig. 5A).

**Fig. S11.**
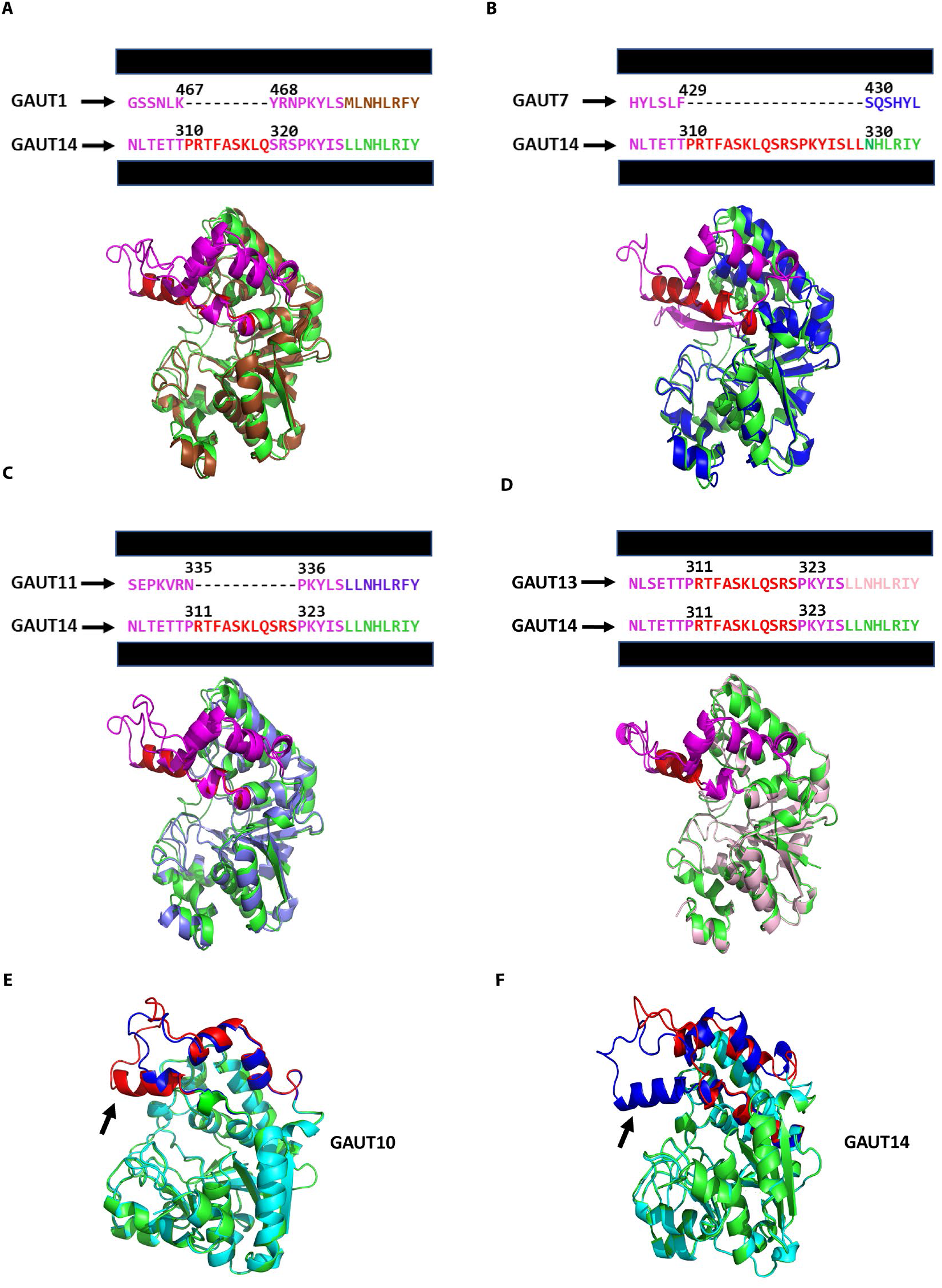
Structural alignment highlights HV1 variability across GAUT family and structural comparison of GAUT10 and GAUT14 wild-type and mutant forms involved in RG-I:HG extension. (A-D) Structural alignments of GAUT14 (green) with GAUT1 (brown, A), GAUT7 (blue, B), GAUT11 (purple, C), and GAUT13 (light pink, D). Alignments were performed across the full GT domain. Only a short sequence window within HV1 is shown above each structure, while the structures display the full HV1 region in the aligned models. The structures highlight variations in the HV1 α-helix (red), with GAUT14 and GAUT13 exhibiting longer HV1 regions containing the *RTFASKLQSRS* motif, whereas GAUT1 and GAUT11 possess short HV1 regions, and GAUT7 even shorter. The HV1 region is shown in purple, except for the α-helix, which is shown in red. The GAUT13 and GAUT14 HV1 regions aligned so well that they are almost superimposable in some places. (E-F) Comparison of AlphaFold-predicted WT and mutant GAUT10 and GAUT14 protein structures. Structures of the HV1 *RTFASKLQSRS* motif-inserted (GAUT10) (E) and motif-deleted (GAUT14) (F) mutants (green) were predicted with AlphaFold. These predicted mutant structures were structurally aligned with their corresponding WT proteins (cyan). The GAUT10 mutant with the inserted motif has a complete predicted HV1 α-helix (black arrow pointing up, red, E) that is absent in the WT (blue). The added α-helix may have provided enough contact surface for the RG-I oligosaccharide to efficiently bind and enabled the mutated GAUT10 to catalyze RG-I:HG extension activity (Fig. 5F). The HV1 region of the GAUT14 mutant protein structure (red), however, lacks the α-helix predicted to be involved in positioning the acceptors in the WT (blue) (black arrow pointing up, F.), and the HV1 region adopts a different fold that could explain why the trifunctional activity is lost in this mutant.

**Fig. S12.**
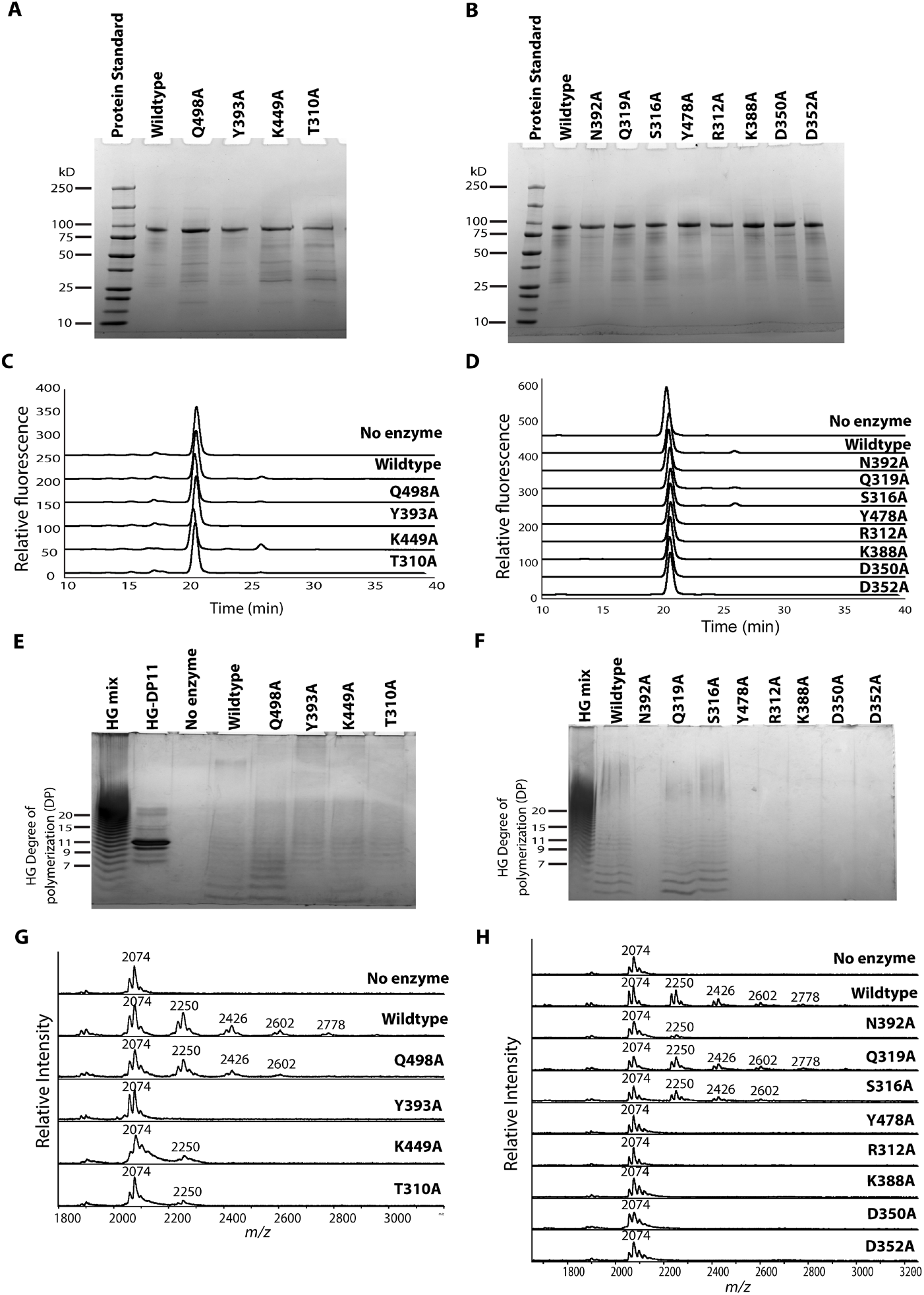
Identification of GAUT14 residues involved in acceptor binding: Single base-pair mutations in GAUT14 amino acids predicted to be involved in acceptor substrate binding (Table S3A-F and S4A-B) and donor substrate coordination (D350A and D352A) (Fig. 4D-G and Fig. S8A-B, F-I) based on molecular modeling were generated. **(A-B)** SDS-PAGE of Ni-NTA^2+^ purified GAUT14 mutants and wildtype (reducing gel). All the mutated proteins electrophoresed close to their expected predicted molecular weight (87.8 kDa). **(C-D)** One-hour qualitative activity assay (using 1 µM enzymes, 1 mM UDP-GalA, 100 µM RG-I-DP10, 50 mM HEPES, and 0.25 mM MnCl2) showing differential effects of site-directed mutation on RG-I:HG extension activity as detected by anion-exchange chromatography with fluorescence detection. Based on the amount of product synthesized, most mutants exhibited knockdown or knockout activity, while S316A and K449A showed slightly increased RG-I:HG extension activity compared to the wild type. **(E-F)** One-hour qualitative activity assay of GAUT14 showing differential effects of site-directed mutagenesis on HG *de novo* synthesis. HG-synthesized products were detected in some mutants but absent in others, as visualized by high-percentage polyacrylamide gel electrophoresis (HP-PAGE). **(G-H)** One-hour qualitative activity assay of GAUT14 mutants showing differential effects on HG acceptor-dependent activity (using 100 µM HG-DP11), detected using MALDI-TOF MS. The mass spectra indicate that all mutants, except Q319A, exhibited knockdown or knockout activity. Mutation of D350 and D352 of the DxD motif knocked out all enzymatic activity, confirming their essential role in donor substrate binding. Y478 and K388 are critical for acceptor binding and catalysis, as mutations of these residues resulted in the complete loss of all three activities.

**Fig. S13.**
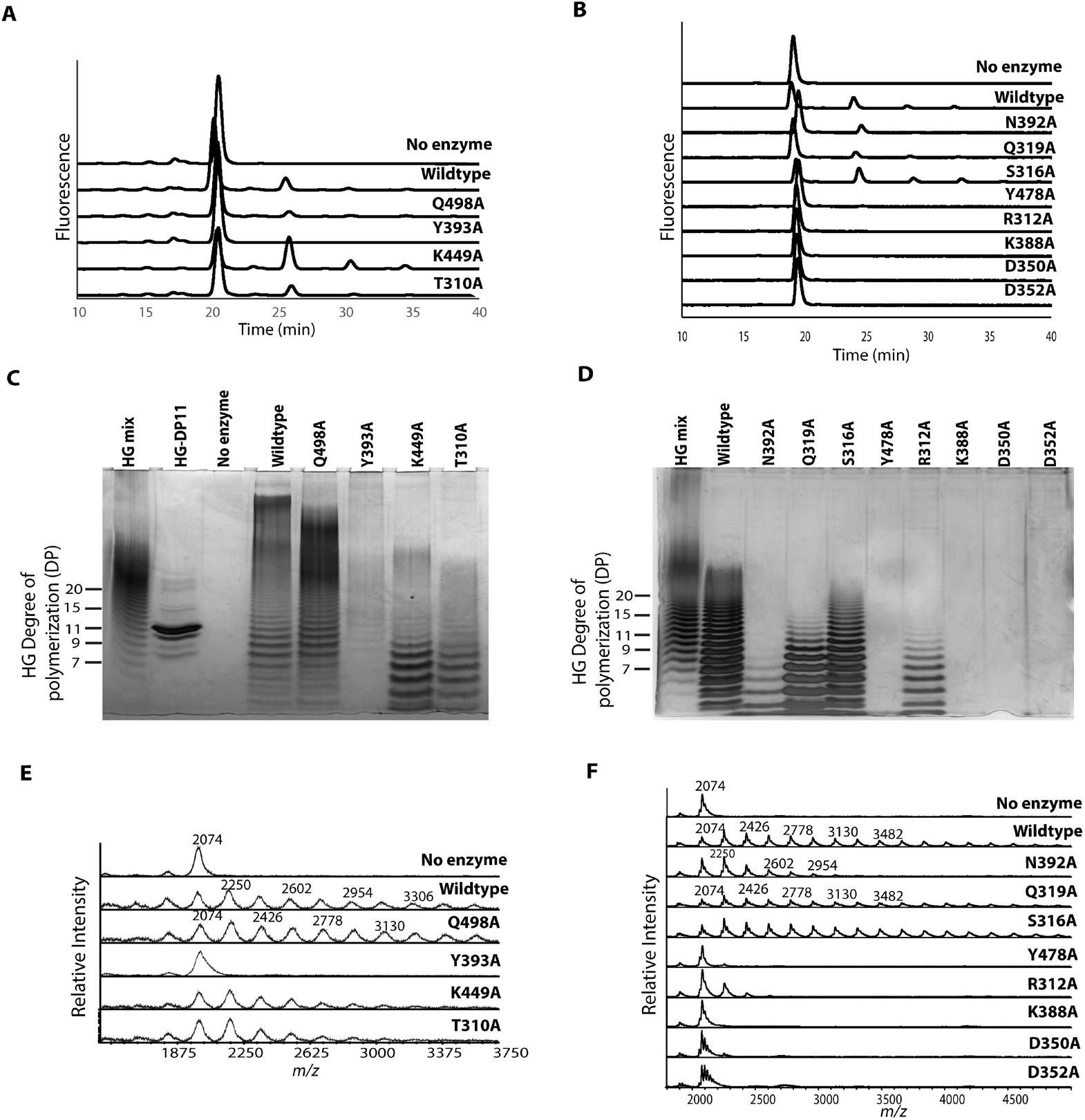
Identification of GAUT14 residues involved in donor and acceptor binding. **(A-B)** Anion-exchange chromatography with fluorescence detection of products produced in 12-hour reactions, showing RG-I:HG extension products synthesized by the GAUT14 mutants versus wild-type GAUT14. Mutants Y478A, R312A, K388A, D350A, and D352A were completely knocked out in RG-I:HG extension activity, while all other mutants, except S316A and K449A, exhibited reduced activity. Notably, mutants S316A and K449A had enhanced RG-I:HG extension activity. **(C-D)** HP-PAGE displaying HG *de novo* synthesis products produced by wild-type GAUT14 and GAUT14 mutants. All mutants tested had knockdown or knockout HG *de novo* synthesis phenotypes (in the case of Y478A, K388A, D350A, and D352A), except for Q498A, which seemingly retained its HG *de novo* synthesis activity compared to the wild type. **(E-F)** MALDI-TOF mass spectrometry analysis of HG acceptor-dependent products synthesized by GAUT14 mutants. Y478A, K388A, Y393A, D350A, and D352A were complete knockouts, while N392A, R312A, K449A, and T310A exhibited reduced HG acceptor-dependent activity.

**Fig. S14.**
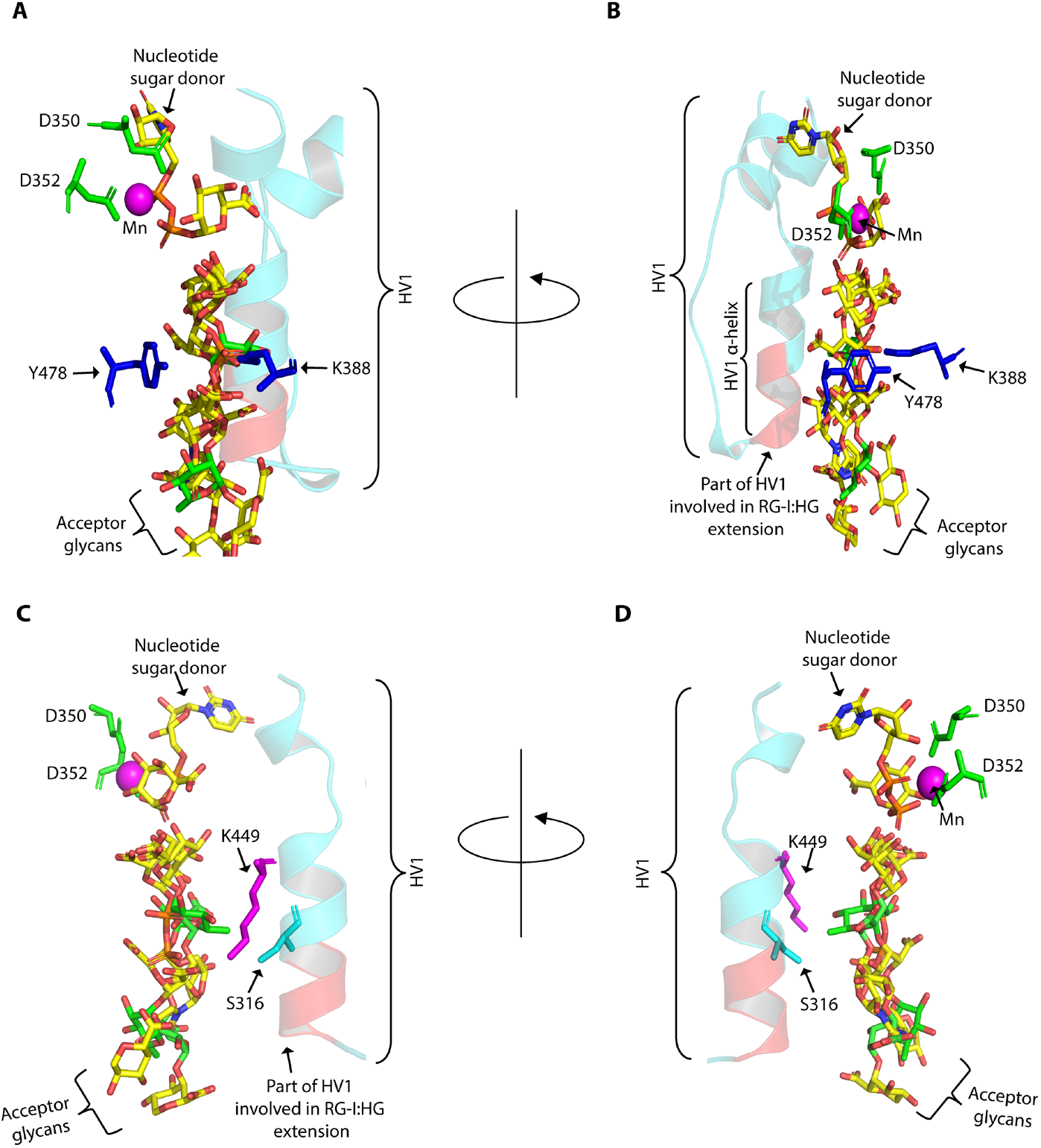
AlphaFold3 protein-glycan model (Model 1) showing residues confirmed via mutagenesis studies (Fig. 5F-I and Fig. S12-13) to be critical in substrate binding. (A-D) D350 and D352 (DxD motif, green sticks) are shown to coordinate the nucleotide sugar donor. The three GAUT14 acceptor substrates are seen in an acceptor-binding groove close to the α-helix of the HV1 region (*RTFASKLQSRS*, cyan), with Y478 and K388 (blue sticks) positioned along the acceptor-binding groove where they likely stabilize or orient the acceptor and growing polysaccharide chain during catalysis (A-B). At the edge of the α-helix, the *PRTFA* residues (red) are required for RG-I:HG extension activity. The positions of amino acids K449 and S316, whose mutation to alanine increased RG-I:HG extension activity, are also shown (C-D). S316 is the fifth amino acid in the *RTFASKLQSRS* HV1 region of the HV1 α-helix while K449 faces the acceptor in the acceptor-binding site.

**Fig. S15.**
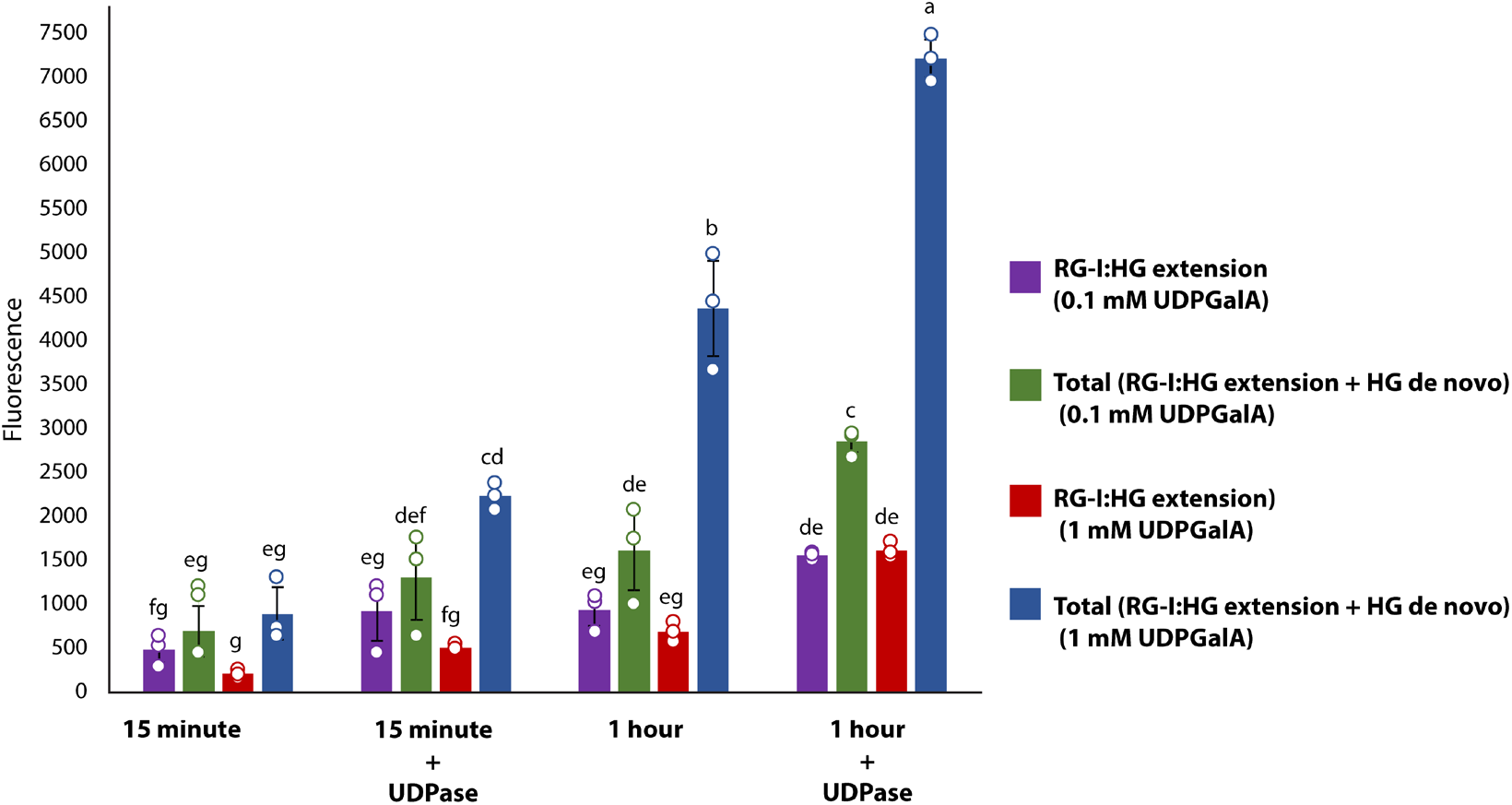
**UDP inhibition of GAUT14 activity is reduced by inclusion of apyrase in the reactions**. Quantification of GAUT14 RGI-HG extension activity in 15 min and 1 h reactions containing 0.1 mM or 1 mM UDP-GalA with or without 20 mU µL^−1^ UDPase (apyrase). Addition of apyrase increased RG-I:HG extension activity and total activity (RG-I:HG extension activity + HG *de novo* synthesis), particularly at 1 mM UDP-GalA and long reaction times, indicating that accumulation of UDP inhibited GAUT14 activity. These results also confirm the lower HG *de novo* synthesis at low 0.1 mM UDP-GalA concentrations and early timepoints (15 min), indicating preferential RG-I:HG extension during the early stages of synthesis. Bars represent mean ± SD of replicate reactions (n=3). Different letters above the bars indicate statistically significant differences between conditions (P < 0.05).

## Supplementary Tables

**Table S1.** Chemical shift assignments of carbohydrate and 2-AB residues in RG-I(G)-DP8+G at pH 2.7 and 7.6. For each group in a residue, the first row shows ^1^H and the second row ^13^C chemical shifts (both in ppm, relative to DSS).

| Sample | Group |  |  |  |  |  | Linkage |
| --- | --- | --- | --- | --- | --- | --- | --- |
| Residue code / ID | 1(a,b) | 2 | 3 | 4 | 5 | 6 |  |
| RG-I(G)-DP8+G |  |  |  |  |  |  |  |
| [pH 2.7] |  |  |  |  |  |  |  |
| A |  |  | 7.09 | 7.53 | 6.98 | 7.64 |  |
| 2-AB | 120.7<sup>a</sup> | 148.5 | 118.1 | 136.5 | 122.1 | 132.0 |  |
| B | 3.67,3.59 | 4.08 | 4.05 | 3.50 | 3.83 | 1.23 | B1—A2 |
| 1N-Rha-ol | 47.0 | 79.3 | 72.4 | 76.4 | 69.8 | 21.3 |  |
| G | 5.03 | 3.89 | 4.12 | 4.45 | 4.97 |  | G1→R2 |
| 4-α-GalpA-(1→2) | 100.4 | 70.4 | 72.2 | 80.5 | 73.0 | 175.5 |  |
| Gb | 5.20 | 3.90 | 4.13 | 4.46 | 4.65 |  | Gb1→B2 |
| 4-α-GalpA-(1→2) | 100.9 | 70.6 | 72.4 | 80.5 | 73.4 | 175.4 |  |
| Gt | 5.08 | 3.86 | 4.09 | 4.47 | 4.89 |  | Gt1→Rt2 |
| 4-α-GalpA-(1→2) | 100.4 | 70.4 | 71.0 | 81.1 | 73.4 | / |  |
| T | 5.09 | 3.74 | 3.94 | 4.33 | 5.03 |  | T1→Gt4 |
| t-α-GalpA-(1→4) | 102.5 | 70.7 | 71.6 | 72.9 | 74.0 | / |  |
| R | 5.21 | 4.11 | 3.86 | 3.43 | 3.66 | 1.25 | R1→G4 |
| 2-α-Rhap-(1→4) | 101.8 | 79.1 | 72.0 | 74.4 | 72.1 | 19.2 |  |
| Rt | 5.22 | 4.15 | / | / | / | / |  |
| 2-α-Rhap-(1→4) | 101.8 | 79.1 | / | / | / | / |  |
| RG-I(G)-DP8+G |  |  |  |  |  |  |  |
| [pH 7.6] |  |  |  |  |  |  |  |
| A |  |  | 6.96 | 7.46 | 6.79 | 7.55 |  |
| 2-AB | 107.0 | 151.0 | 115.8 | 136.3 | 119.3 | 131.9 |  |
| B | 3.61,3.51 | 4.03 | 4.00 | 3.49 | 3.82 | 1.24 | B1—A2 |
| 1N-Rha-ol | 44.6 | 78.5 | 72.1 | 76.3 | 69.9 | 21.4 |  |
| G | 4.99 | 3.91 | 4.11 | 4.40 | 4.65 |  | G1→R2 |
| 4-α-GalpA-(1→2) | 100.1 | 70.6 | 73.0 | 80.0 | 74.0 | / |  |
| Gb | 5.12 | 3.88 | 4.07 | 4.40 | 4.35 |  | Gb1→B2 |
| 4-α-GalpA-(1→2) | 100.4 | 70.7 | 73.2 | 80.0 | 74.3 | 177.2 |  |
| Gt | 5.03 | 3.84 | 4.06 | 4.43 | 4.69 |  | Gt1→Rt2 |
| 4-α-GalpA-(1→2) | 100.2 | 70.6 | 71.5 | 80.9 | 73.9 | / |  |
| T | 5.09 | 3.73 | 3.92 | 4.27 | 4.75 |  | T1→Gt4<sup>b</sup> |
| t-α-GalpA-(1→4) | / | 71.1 | 72.3 | 73.6 | 75.0 | / |  |
| R | 5.25 | 4.10 | 3.88 | 3.40 | 3.78 | 1.25 | R1→G4 |
| 2-α-Rhap-(1→4) | 101.2 | 78.7 | 72.1 | 74.6 | 71.7 | 19.3 |  |
| Rt | / | 4.13 | / | / | / | / |  |
| 2-α-Rhap-(1→4) | / | 78.9 | / | / | / | / |  |

<sup>a</sup> <sup>13</sup>C chemical shift of the 2-AB amide carbonyl is 176.2 ppm.
<sup>b</sup> Very weak NOE at pH 7.6.
/, not determined

**Table S2:** Metrics supporting GAUT14 as a dimer rather than a trimer. Analysis of the GAUT14 AlphaFold-predicted dimer and trimer structures suggests that GAUT14 is a dimer. This is supported by the following metrics, all of which meet their respective cutoffs for a multimer: IpTM + pTM (> 0.7), ratio of interface area (> 4%), interface confidence level (> 80%), ΔG kcal/mol (< -20 kcal/mol), and ΔG p-value (< 0.3). The cutoffs were determined based on the analysis of multiple AlphaFold-predicted structures that have been previously shown to exist as multimers, such as GAUT1:GAUT7.

| Metrics and cutoffs<br>(in round brackets) | GAUT14<br>Dimer | GAUT14<br>Trimer | GAUT1:GAUT7<br>Dimer |
| --- | --- | --- | --- |
| lptm + ptm<br>(> 0.7) | 0.78 | 0.43 | 0.71 |
| Ratio of Interface Area<br>(> 4 %) | 4.7 | 3 | 4.7 |
| Interface confidence<br>level<br>(> 80 %) | 81 | 71.5 | 83 |
| Δg kcal/mol<br>(< -20kcal/mol) | -56.1 | -20.7 | -40.9 |
| Δg p-value<br>(< 0.3) | 0.021 | 0.466 | 0.24 |

**Table S3:** MM/GBSA binding free energy calculation of GAUT14 residue interactions with UDP-GalA, RG-I-DP5, and HG-DP5 acceptors. The AlphaFold3 protein-glycan model (Model 1) from Fig. 4D-G was subjected to molecular dynamics (MD) simulation and the trajectories of the structure with its bound substrate during simulation were subjected to Molecular Mechanics/Generalized Born Surface Area (MM/GBSA) per-residue energy analysis to estimate the contribution of individual amino acids to glycan binding (black; A,C,E). This calculation was also carried out for the trajectories of the docking/MD simulated model (Model 2, Fig. S8 F-I) (blue; B,D,F). The table shows the total free energy contributions for each residue (kcal/mol). Negative total ΔG values indicate favorable contributions to binding. Residues with ΔG ≤ −2.0 kcal/mol were considered strong contributors to substrate binding; those between −1.0 and −2.0 kcal/mol were classified as moderate contributors, and residues with ΔG < 1.0 kcal/mol were considered weak contributors (Not included in the Table).

| RG-I Interacting residues | vdW | Electro-statics | Polar | Nonpolar | Total ($\Delta G$) |
| --- | --- | --- | --- | --- | --- |
| LYS 388 | -0.1 | -72.3 | 57 | -1.0 | -16 |
| ASN 496 | -0.5 | -8.5 | 3.2 | -0.4 | -6.2 |
| ASN 392 | -0.6 | -6.7 | 2.1 | -0.8 | -6.1 |
| SER 316 | -1.1 | -2.0 | 0.8 | -0.9 | -3.3 |
| LYS 449 | -0.6 | -37.5 | 36 | -0.6 | -2.5 |
| THR 313 | 0.88 | -0.2 | -0.8 | -0.7 | -2.1 |
| TYR 393 | -0.4 | -2.7 | 1.4 | -0.3 | -2.0 |
| TYR 478 | -0.9 | -0.9 | 0.5 | -0.5 | -1.9 |
| ARG 391 | -0.5 | -31.9 | 31.1 | -0.4 | -1.7 |
| LEU 450 | -0.9 | -0.3 | 0.2 | -0.7 | -1.7 |
| ARG 312 | -0.4 | -27.9 | 27 | -0.3 | -1.2 |

B
| RG-I Interacting residues | vdW | Electro-statics | Polar | Nonpolar | Total ($\Delta G$) |
| --- | --- | --- | --- | --- | --- |
| ARG 312 | -1.8 | -58.3 | 51.6 | -2.0 | -10.6 |
| LYS 388 | -0.9 | -52.1 | 44.0 | -1.5 | -10.5 |
| TYR 478 | -1.2 | -3.7 | 1.6 | -1.2 | -4.6 |
| ASN 392 | -1.1 | -2.2 | 0.3 | -0.9 | -3.9 |
| SER 316 | -0.8 | -0.3 | -0.5 | -0.8 | -2.4 |
| GLN 319 | -0.6 | -3.8 | 2.8 | -0.7 | -2.3 |
| LYS 449 | -0.5 | -31.7 | 30.7 | -0.5 | -2.0 |
| THR 313 | -0.4 | -0.6 | -0.6 | -0.4 | -2.0 |
| ASN 496 | -0.4 | -2.0 | 0.8 | -0.3 | -1.9 |

C
| HG Interacting residues | vdW | Electro-statics | Polar | Nonpolar | Total ($\Delta G$) |
| --- | --- | --- | --- | --- | --- |
| ARG 312 | -0.9 | -73.4 | 67 | -1.2 | -7.7 |
| LYS 388 | -0.1 | -70.7 | 66.2 | -0.5 | -5.1 |
| TYR 478 | -1.1 | -0.2 | -0.1 | -1.0 | -2.6 |
| ASN 496 | -0.4 | -6.8 | 5.3 | -0.3 | -2.2 |
| ASN 392 | -0.4 | -2.9 | 1.6 | -0.4 | -2.0 |
| GLN 498 | -0.6 | -4.4 | 3.5 | -0.5 | -1.9 |
| GLY 497 | -0.5 | -3.6 | 3.3 | -0.4 | -1.3 |
| LYS 449 | -0.1 | -55.0 | 54.0 | -0.1 | -1.2 |
| GLN 319 | -0.2 | -1.1 | 0.4 | -0.2 | -1.2 |

D
| HG interacting residues | vdW | Electro-statics | Polar | Nonpolar | Total ($\Delta G$) |
| --- | --- | --- | --- | --- | --- |
| LYS 388 | -0.1 | -82.6 | 72.4 | -0.9 | -11.3 |
| ARG 312 | -1.1 | -72.5 | 68.5 | -1.1 | -6.2 |
| LYS 449 | -0.2 | -63.0 | 60.2 | -0.4 | -3.4 |
| TYR 478 | -1.4 | 0.6 | -0.7 | -1.2 | -2.7 |
| TRP 384 | -0.6 | 0.2 | -1.3 | -1.2 | -1.6 |
| ASN 496 | -0.3 | -4.5 | 3.7 | -0.3 | -1.5 |
| TYR 393 | -0.4 | -1.1 | 0.5 | -0.5 | -1.4 |
| THR 310 | -0.1 | -1.7 | 1.4 | -0.8 | -1.2 |

E
| UDP-GalA<br>Interacting<br>residues | vdW | Electro-statics | Polar | Nonpolar | Total<br>(ΔG) |
| --- | --- | --- | --- | --- | --- |
| ARG 312 | -0.8 | -4.0 | 1.4 | -0.8 | -4.2 |
| TYR 478 | -1.4 | -0.6 | 0.4 | -1.1 | -2.6 |
| LYS 388 | -0.2 | -0.8 | -0.3 | -0.3 | -1.7 |
| GLN 498 | -0.4 | -0.6 | 0.04 | -0.4 | -1.4 |
| ASN 392 | -0.3 | 0.2 | -0.04 | -0.2 | -0.9 |
| GLY 497 | -0.3 | -0.2 | -0.05 | -0.3 | -0.9 |

F
| UDP-GalA<br>Interacting<br>residues | vdW | Electro-statics | Polar | Nonpolar | Total<br>(ΔG) |
| --- | --- | --- | --- | --- | --- |
| LYS 449 | -0.3 | -59.0 | 52.7 | -0.6 | -7.2 |
| ARG 312 | -1.5 | -34.3 | 31.4 | -1.2 | -5.7 |
| LYS 388 | 0.0 | -50.8 | 46.3 | -0.3 | -4.8 |
| TYR 478 | -0.8 | -3.5 | 2.1 | -0.9 | -3.1 |
| THR 313 | -0.6 | 0.4 | -0.9 | -0.4 | -1.4 |
| TRP 384 | -1.0 | 2.6 | -2.4 | -0.7 | -1.4 |
| GLN 498 | -0.3 | -4.8 | 4.0 | -0.2 | -1.2 |
| SER 316 | -0.5 | -1.5 | 1.3 | -0.4 | -1.1 |

**Table S4:** Amino acid residues predicted to contribute to GAUT14 acceptor binding based on hydrogen bond occupancy from Molecular Dynamics simulation. This table presents the top predicted amino acid residues involved in binding of all three acceptors to GAUT14, based on hydrogen-bond occupancy analysis. The occupancy (%) represents the percentage of time during the MD simulation that a hydrogen bond was present or formed between a given amino acid residue and each acceptor substrate. The occupancy was calculated for the simulated AlphaFold3 protein-glycan model (black; A) and for the docking/MD simulated model (blue; B). Values represent mean ± standard deviation (n = 3 independent experiments).

| Amino acid residues | UDP-GalA (%) | HG (%) | RG-I (%) |
| --- | --- | --- | --- |
| TYR478 | | 5.5 ± 3.5 | 14.9 ± 2 |
| LYS388 | 3.7 ± 7.3 | 6.7.7.2 | 50.1 ± 14.1 |
| GLN498 | 8.6 ± 3.9 | 8.9 ± 6.4 | |
| GLN319 | 2.2 | 7.2 ± 1.8 | 19.3 ± 6.4 |
| TYR393 | | 3.7 | 28.3 ± 5.5 |
| LYS449 |  | 3.3 | 2.0 |
| ARG312 | 15.1 ± 7.2 | 28.1 ± 9.5 | |
| SER316 | | 3.7 | 9.4 ± 4.2 |
| ASN392 | 3.9 ± 1.9 | 12.1 ± 1.3 | 69.5 ± 23.2 |
| THR310 |  | 2.2 |  |
| TYR325 | | 16.8 ± 2.2 | 2.7 ± 1.1 |
| ASN496 | 2.0 ± 4.4 | 6.1 | 30.5 ± 16.2 |
| THR313 |  |  | 5.8 |

B
| Amino acid residues | UDP-GalA (%) | HG (%) | RG-I (%) |
| --- | --- | --- | --- |
| TYR478 | 34.9 ± 14.9 | 20.2 ± 9.7 | 15.4 ± 6.4 |
| LYS388 | 42.6 ± 3.7 | 46.7 ± 5.2 | 79 ± 8.8 |
| GLN498 | 66.8 ± 2.2 | | |
| GLN319 | 2.5 ± 1.5 | | 24 |
| TRP384 | 2.4 |  |  |
| TYR393 | 44.3 ± 11.4 | 1.8 | 77.9 ± 31 |
| LYS449 | 19.7 ± 18.1 | 32.7 ± 19 | 23.4 ± 15.6 |
| ARG312 | 58.9 ± 6.6 | 100 | 57.5 ± 7.4 |
| GLN479 | 2.2 ± 2.1 | 2.6 ± 2.7 | |
| SER316 | 7.2 ± 2.8 | 7.2 ± 3.2 | 6.1 ± 1.5 |
| LYS160 | 2.6 |  |  |
| ASN392 | 31.1 | 17.1 | 26.7 |
| LYS144 |  | 6.9 |  |
| THR310 | | 40 | 39.6 ± 7.6 |

**Table S5:**
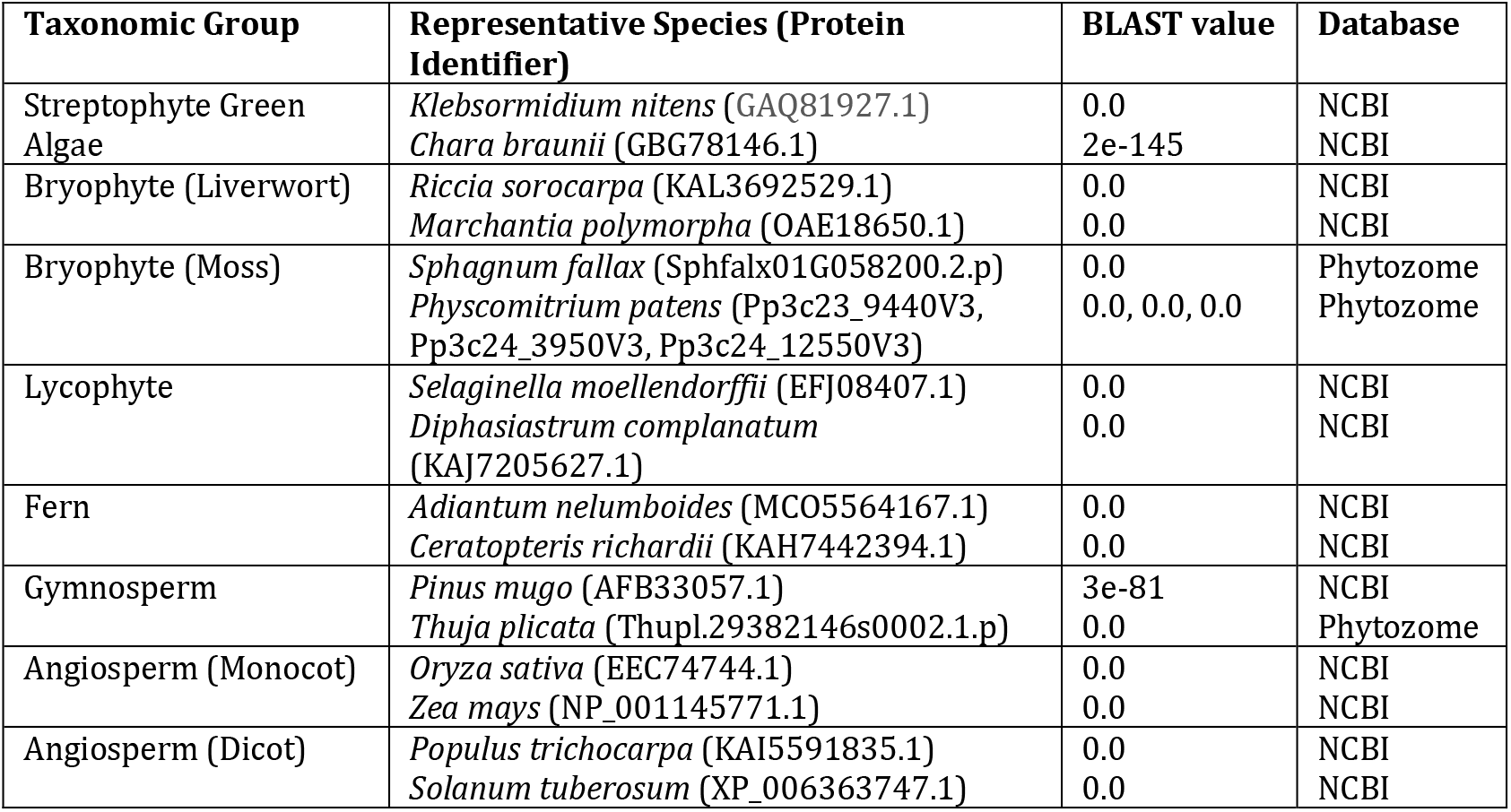
Identification of GAUT13/14 homologs across major plant lineages. Identification of GAUT13/14 homologs across major plant lineages. NCBI and Phytozome PBLAST searches showed that GAUT13 and GAUT14 homologs are present from streptophytic green algae to flowering plants (angiosperms). The table lists the selected taxa, representative species (and their protein identifier), PBLAST value, and source databases. An E-value of 0.0 indicates an extremely high sequence similarity, where the calculated probability is so low that it rounds down to 0.0.

**Table S6:**
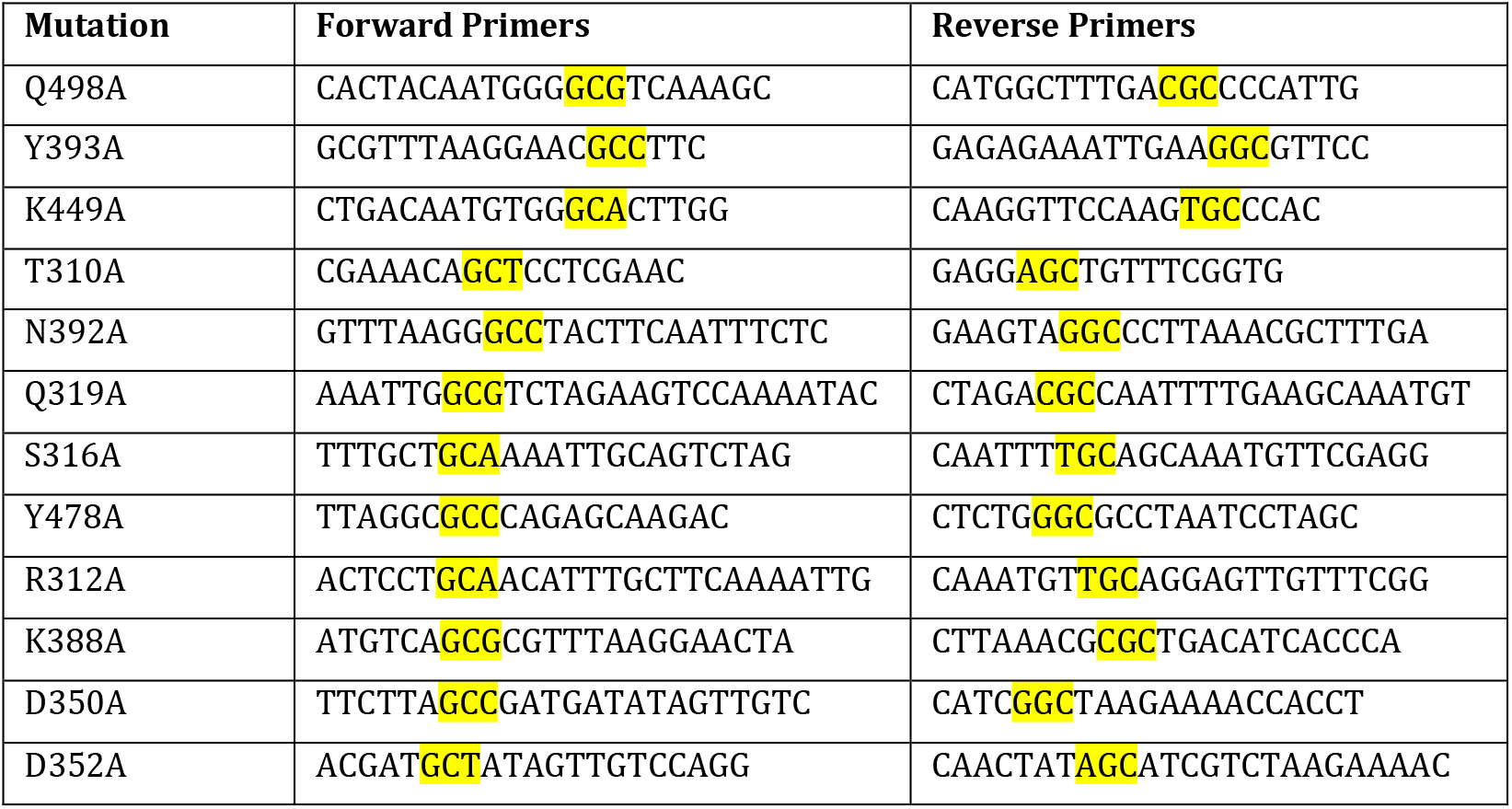
Forward and reverse primers used for GAUT14 site-directed mutagenesis. These primers were designed to introduce single amino-acid substitutions into the GAUT14 coding sequence to identify amino acid residues involved in GAUT14 substrate binding. The mutations were introduced into GAUT14 using a GAUT14:pDONR^TM^221 construct by PCR-based site-directed mutagenesis. Yellow highlight indicates the mutated codons on primers introduced to generate alanine mutations at the indicated residues.

